# Nutritional signals rapidly activate oligodendrocyte differentiation in the adult hypothalamic median eminence

**DOI:** 10.1101/751198

**Authors:** Sara Kohnke, Brian Lam, Sophie Buller, Chao Zhao, Danae Nuzzaci, John Tadross, Staffan Holmqvist, Katherine Ridley, Hannah Hathaway, Wendy Macklin, Giles SH Yeo, Robin JM Franklin, David H Rowitch, Clemence Blouet

**Affiliations:** MRC Metabolic Diseases Unit, University of Cambridge Metabolic Research Laboratories, WT-MRC Institute of Metabolic Science, University of Cambridge, Cambridge CB2 OQQ, UK; Wellcome -MRC Cambridge Stem Cell Institute and department of Clinical Neurosciences, -University of Cambridge, Cambridge, UK; Department of Cell & Developmental Biology, University of Colorado School of Medicine, Aurora, CO, USA; Program in Neuroscience, University of Colorado School of Medicine, Aurora, CO, USA; Department of Paediatrics and Wellcome-MRC Cambridge Stem Cell Institute, University of Cambridge, Cambridge, UK

**Author notes:** Dr Clémence Blouet, Metabolic Research laboratories, IMS level 4, Box 289, Addenbrookes Hospital, Hills Road, Cambridge, CB2 0QQ, UK.

## Abstract

The mediobasal hypothalamus (arcuate nucleus - ARC - and median eminence - ME -) controls energy balance, growth and fertility through its ability to integrate neuronal, nutritional and hormonal signals and coordinate the behavioural, neuroendocrine and metabolic responses required for these functions. While our understanding of the neural circuits downstream from ARC neurons is rapidly progressing, little is known about the function of other cell types. Here we describe an unexpected role for oligodendrocytes (OL) of the ME in monitoring nutritional signals. We show that refeeding following an overnight fast rapidly activates oligodendrocyte differentiation and the production of new OL in the ME specifically. No changes in myelination were measured in this time-frame. However, refeeding changed the expression of OL-derived extracellular matrix proteins decorin and tenascin-R, with consistent changes in the density of local perineuronal nets. Last, we show that OLs use mTORC1 signalling, a pathway required for OL differentiation, to survey energy and protein availability, specifically in the ME. We conclude that new oligodendrocytes formed in the ME in response to nutritional signals control the access of circulating metabolic cues to ARC interoceptive neurons.

## Introduction

The median eminence (ME) of the mediobasal hypothalamus (MBH) is a bidirectional gateway between the hypothalamus and the periphery with diverse roles in mammalian physiology including energy homeostasis and neuroendocrine control of growth, reproduction, lactation, stress and the thyroid axis. The ME vasculature lacks a blood-brain barrier (BBB), allowing axons of hypothalamic neuroendocrine neurons to access a BBB-free area when entering the ME, and release hypothalamic-releasing hormones into fenestrated capillaries that carry blood to the pituitary. The ME fenestrated endothelium also allows circulating signals to freely diffuse into the ME and adjacent arcuate nucleus of the hypothalamus (ARC), rich in neurons critical to appetite regulation and energy balance, thus giving local neurons privileged access to peripheral signals.

Tanycytes of the ME have been proposed to replace the lacking endothelial BBB and control the diffusion of blood-borne signals from the ME to the adjacent ventricular space through their well-organized tight junctions (ME-CSF barrier)^1^. The ME-ARC barrier is functionally equally critical, but its structural components remain unknown^2^. In addition, ME tanycytes regulate the release of hypothalamic releasing hormones into the ME portal system^3,4^. Little is known about how other glial cell types contribute to these functions.

Emerging evidence indicates that the ME is highly plastic and rapidly responds to hormonal and nutritional signals ^5,6^. **T**his could relate to its unique unbuffered access to blood-borne signals, specifically exposing the ME to acute peripheral changes. Hormonal and nutritional sensing in the ME initiates rapid structural remodelling, leading to changes in the ARC territory sitting outside the BBB and exposed to peripheral signals^7^. ME tanycytes have been proposed to mediate the nutritional regulation of the ME-CSF barrier^7^, but the mechanisms mediating the plasticity of the ME-ARC barrier in response to nutritional cues remain to be determined.

Here we report a novel function for oligodendrocytes (OLs) of the ME in response to nutritional signals. We found that the adult ME is rich in newly-formed oligodendrocytes. These, together with mature OLs, are concentrated at the ME-ARC barrier. We found that nutritional signals rapidly activate OL differentiation specifically in the ME and regulate the formation of perineuronal nets. Collectively, these studies reveal an unsuspected plasticity of oligodendrocytes of the ME in response to nutritional signals with functional consequences contributing to the regulation of neuronal exposure to blood-borne signals.

## Results

### Single cell transcriptomic analysis of median eminence cell populations

We used single-cell RNA sequencing to build a high resolution cellular and molecular characterization of the adult mouse median eminence (ME) and understand the transcriptional changes occurring during a nutritional transition from the fasted to the fed state (Figure 1a, Supplementary Figure 1a-c). We mapped each of the 5,982 sequenced cells onto a t-distributed stochastic neighbor embedding (tSNE) plot based on gene expression and identified 9 distinct cell clusters using K-means clustering (Figure 1b, Supplementary Figure 1d). We obtained the unique transcriptional signature of each cluster (Figure 1c, Supplementary Table 3) and a defining gene for each cluster, as follows: tanycytes (*Rax*^+^), astrocytes (*Agt*^+^), oligodendrocytes (type 1, *Ermn*^+^), microglial cells (*C1qc*^+^), neurons (*Snhg11*^+^), oligodendrocytes (type 2, *Cd9*^+^), vascular and leptomeningeal cells (VLMC, *Dcn*^+^), ependymocytes (*Elof1*^+^), and endothelial cells (*Itm2a*^+^) (Figure 1d). We found that neurons represented only 9.1% of the total cell population of the ME, most of which were GABAergic (Supplementary Figure 1e). While other cell types did not separate into distinct clusters on the tSNE plot, oligodendrocytes showed 2 clusters, suggesting diversity in this cell lineage (Figure 1B).

**Figure 1.**
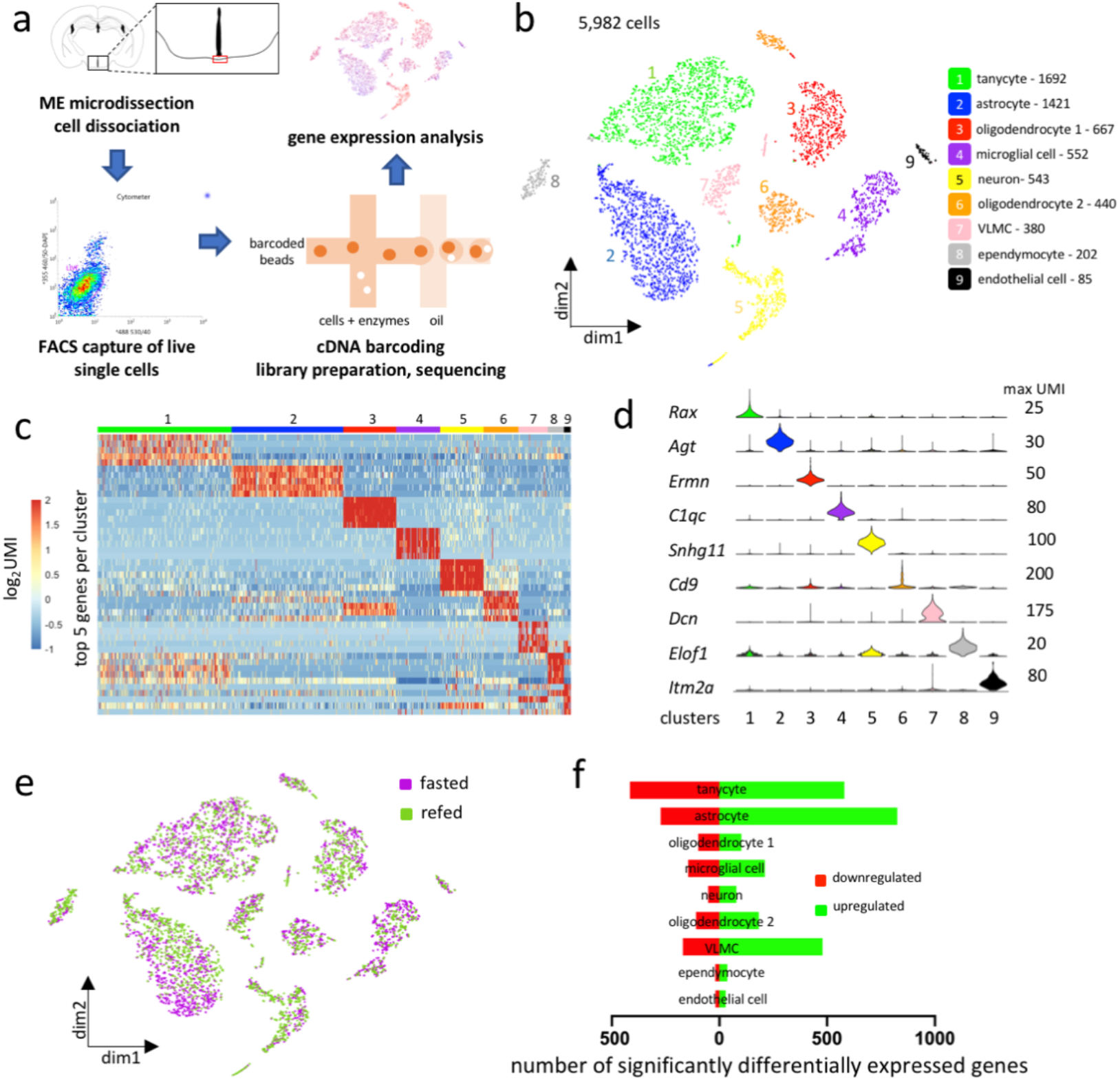
Nine cell types are present in the ME. (a) Workflow for single-cell RNA sequencing experiment (b) t-distributed stochastic neighbor embedding (tSNE) plot demonstrating 9 groups of ME cells). VLMC = vascular and leptomeningeal cells (c) Heatmap of log_2_UMI counts per cell for the top 5 differentially-expressed genes per cluster (d) Violin plot showing UMI count distribution of one defining gene per cluster (variable scales per gene) (e) Cell sample IDs mapped on tSNE plot (n = 5 per condition) (f) Number of genes significantly different between fasted and refed conditions per cluster (*p* < 0.05 and FDR < 0.25)

Cells from the fasted and refed samples were generally well-mixed across all clusters, indicating that fasted and refed conditions do not change gross identities of cells (Figure 1e). We obtained the list of genes significantly differentially expressed between the two nutritional conditions in each of the 9 clusters (Figure 1f, Supplementary Table 4). Tanycytes, astrocytes, and VLMC were the most nutritionally responsive cell types, whereas neurons had relatively few differentially expressed genes between the fasted and refed conditions. A total of 495 genes were differentially expressed in OLs, revealing a previously unknown rapid nutritional regulation of this cell type.

### Gene profiling suggest three states of OL lineage cells in the ME

The evidence for molecular diversity and nutritional regulation of ME OLs prompted us to further study this cellular population. From the previous nine clusters, we reclustered cells present in cluster 3 (oligodendrocyte 1) and cluster 6 (oligodendrocyte 2), which revealed 3 subtypes of OLs with distinct molecular signatures (Figure 2a-c, Supplementary Figure 2a, Supplementary Table 5).

**Figure 2.**
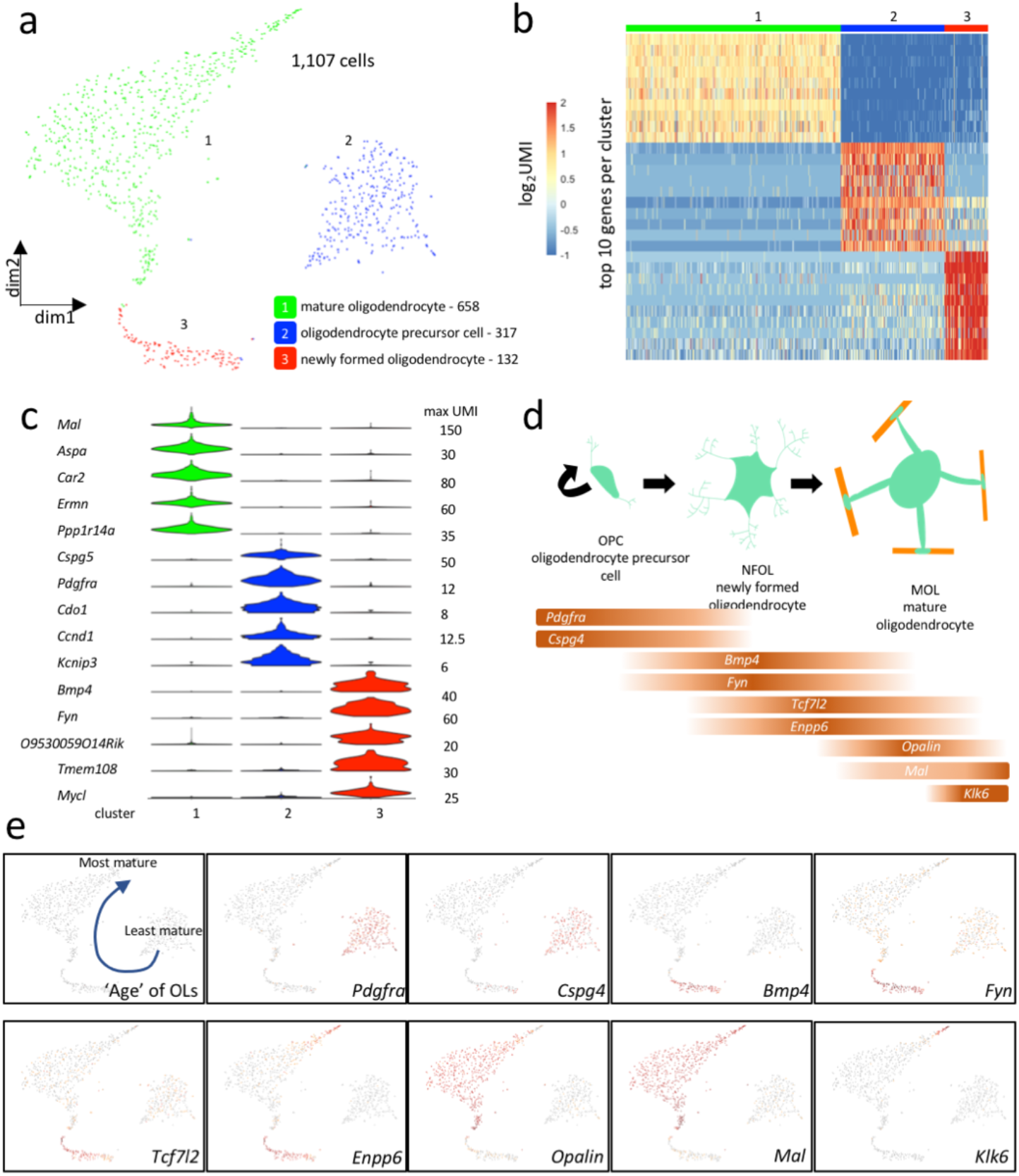
Three gene expression states of OL lineage cells are present in the ME. (a) OLs segregate into 3 clusters on a t-distributed stochastic neighbor embedding (tSNE) plot (b) Heatmap of log_2_UMI counts per cell for the top 10 differentially-expressed genes per cluster (c) Violin plot showing UMI count distribution of the top 5 genes per cluster (variable scales per gene). (d) Schematic of oligodendrocyte differentiation, and expression of validated stage markers in our OL database (e) tSNE plots showing expression of validated oligodendrocyte markers in current dataset. Red = high expression, grey = low expression

Established molecular markers of OL subtypes allowed us to characterize cells in OL cluster 2 as oligodendrocyte progenitor cells (OPCs, express *Pdgfra* and *Cspg4*), an immature cell type that differentiates into OLs ^8,9^ (Figure 2b-e). Cells in cluster 3 expressed genes associated with differentiating OLs. Here we named them newly formed oligodendrocytes (NFOLs) as they express *Bmp4*, *Fyn, Tcf7l2,* and *Enpp6* ^8,10^ (Figure 2c-e, Supplementary Figure 2b). We named cells in cluster 1 mature oligodendrocytes (MOLs) as they expressed *Opalin*, *Mal,* and *Klk6* ^8^ (Figure 2c-e, Supplementary Figure 2b). The distribution of cells expressing stage-specific genes indicates that the coordinates of OLs in the tSNE plot reflect the cell’s developmental ‘age,’ which follows a curve in the plot (Figure 2e, Supplementary Figure 2c).

### The NFOL and MOL populations are concentrated in the dorsal part of the murine and human ME

We used RNAscope single-molecule fluorescence *in situ* hybridization (FISH) to map the density and neuroanatomical distribution of OL lineage subtypes in the murine ME and ARC, and used the corpus callosum (CC) as a reference tissue (Figure 3a, Supplementary Figure 3a-c). We found that OPCs (*Pdgfra^+^*/Sox10^+^ cells) are evenly distributed throughout the ME and ARC (Figure 3a), with densities of OPCs in both structures similar to that measured in the CC (Figure 3g, Supplementary Fig 3b). In contrast, the density NFOLs (*Bmp4*^+^and/or *Tcf7l2^+^*/Sox10^+^) and MOLs (*Plp1*^+^ and Sox10^+^) was more than 3-times higher in the ME than in the ARC (Figure 3a, g). Strikingly, NFOLs and MOLs were almost absent from the ARC and occupied exclusively the dorsal portion of the ME extending bilaterally to the ventral base of the ARC (Figure 3a).

**Figure 3.**
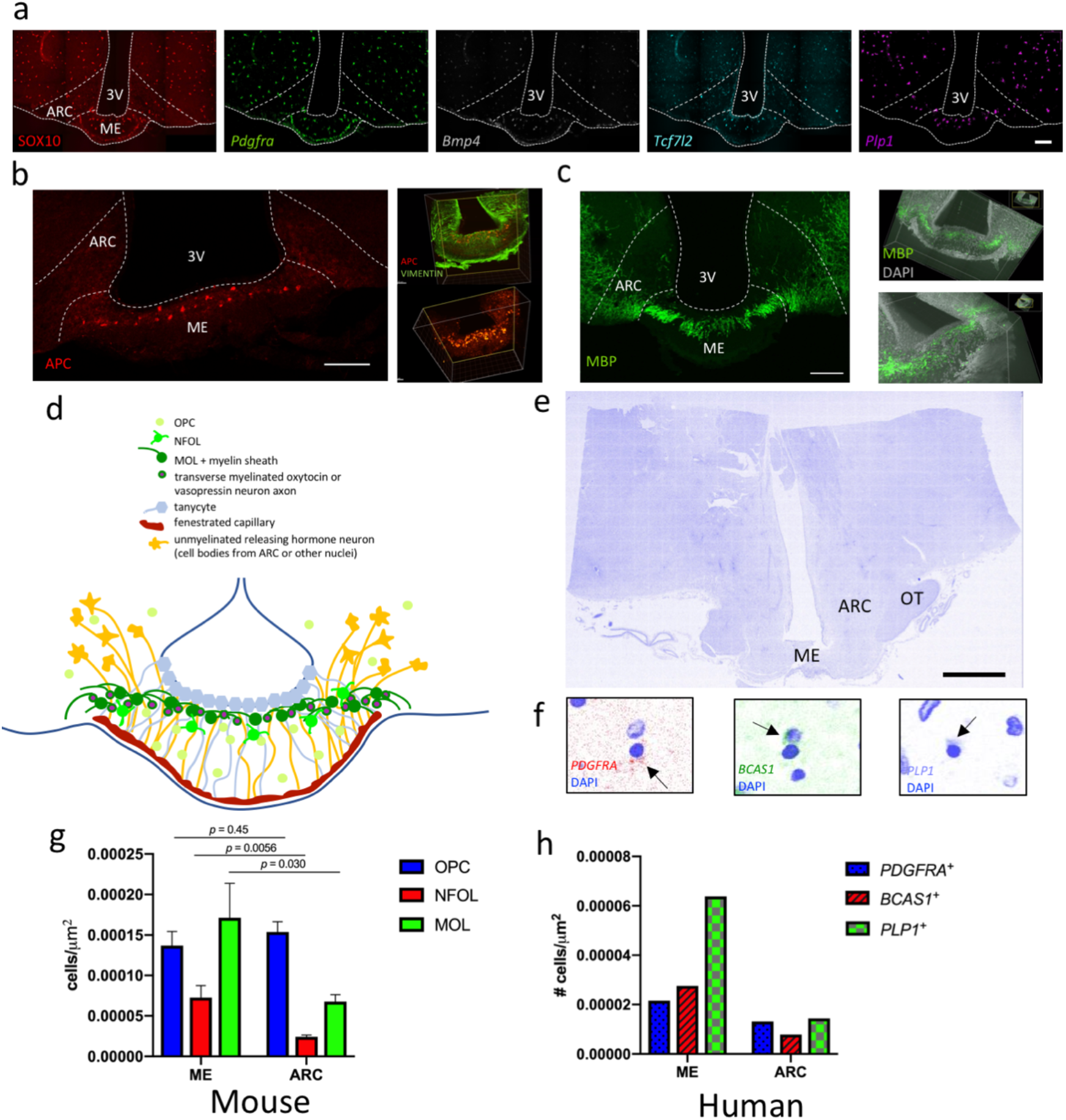
The NFOL and MOL populations are concentrated in the dorsal part of the murine and human ME. (a) FISH combined to immunohistochemistry for detection of OL subtype marker gene expression in mouse brain: red = Sox10 (IHC, pan-OL marker), green = *Pdgfra* (OPC marker), grey = *Bmp4* (NFOL marker), blue = *Tcf7l2* (NFOL marker), purple = *Plp1* (MOL marker). Scale bar = 100 μm. (b) APC protein (MOL marker, red) in thin coronal sections (left, scale bar = 100 μm) and APC and vimentin (tanycyte marker, green) in thick cleared ME tissue (right, screenshots from Video 1). (c) MBP protein (myelin marker, green) in thin coronal sections (left, scale bar = 100 μm) and in thick cleared ME tissue (right, screenshots from Video 2). (d) Schematic of different cell types of the ME (e) DAPI stain in human hypothalamus. ME = median eminence, ARC = arcuate nucleus, OT = optic tract, scale bar = 3.75 mm. (f) Probes to PDGFRA, BCAS1, and PLP1 label OPCs, NFOLs, and MOLs of the human hypothalamus, respectively. Arrowheads indicate probe labelling. (g) Measurement of densities of OPCs (*Pdgfra*^+^/Sox10^+^), NFOLs (*Bmp4*^+^ and or *Tcf7l2*^+^/Sox10^+^), and MOLs (*Plp1*^+^/Sox10^+^) in the mouse ME and ARC. Error bars depict mean ± SEM, p values determined with a two-sided Students t-test (ME n = 6 animals, ARC n = 9 animals) (h) Measurement of densities of OPCs (*PDGFRA^+^),* NFOLs (BCAS1^+^), and MOLs (*PLP1*^+^) in the human ME and ARC. (n = 1 human)

To gain a better understanding of how postmitotic OLs are positioned in the ME, we used tissue clearing and immunolabelling for APC (CC1 clone), used here as a marker of MOLs^11^. Consistent with our findings with FISH, we found that APC^+^ cells populate exclusively the dorsal portion of the ME, immediately below the layer of vimentin^+^ tanycytes (Figure 3b, Supplementary video 1). Labelling of myelin basic protein (MBP) revealed the presence of dense myelin fibers in the dorsal ME and under the base of the ARC, but absence of myelin in the ARC itself (Figure 3c, Supplementary video 2). Along with knowledge of how neurons and their axons and fenestrated capillaries are positioned in the ME^4,7^, we were able to construct a model that showcases the specific and discrete localization of NFOLS and MOLs in the ME (Figure 3d).

We then used hypothalamic brain sections from an adult human donor to characterise the neuroanatomical distribution of OL subtypes in the human mediobasal hypothalamus and investigate the presence of NFOLs there. In the adult human brain, BCAS1 is marker of NFOLs^12^. We used FISH to visualize *PDGFRA* (OPCs), *BCAS1* (NFOLs), and *PLP1* (MOLs), and compared the densities of cells expressing the genes in the ME and ARC of human tissue (Figure 3e, f, h). These experiments revealed the presence of NFOLs, indicating the continuous formation of new OLs in the adult human ME. As in the mouse mediobasal hypothalamus, we found that OPCs were present at similar densities in the ME and ARC, whereas the ME contains 3.5 times more NFOLs and 4.4 times more MOLS than the ARC (Figure 3h).

### Nutritional signals rapidly regulate the OL transcriptome in the adult ME

To begin to characterise the functional consequences of the fast-refeed stimulus on oligodendrocyte populations, we looked at differentially-expressed genes (DEGs) in fasted versus refed ME samples in the 3 oligodendrocyte clusters (Figure 4a, Supplementary Table 6). In all 3 clusters, genes regulating OL lineage progression (OPC migration, proliferation, cell cycle exit and differentiation) represented a large proportion of DEGs, as follows.

**Figure 4.**
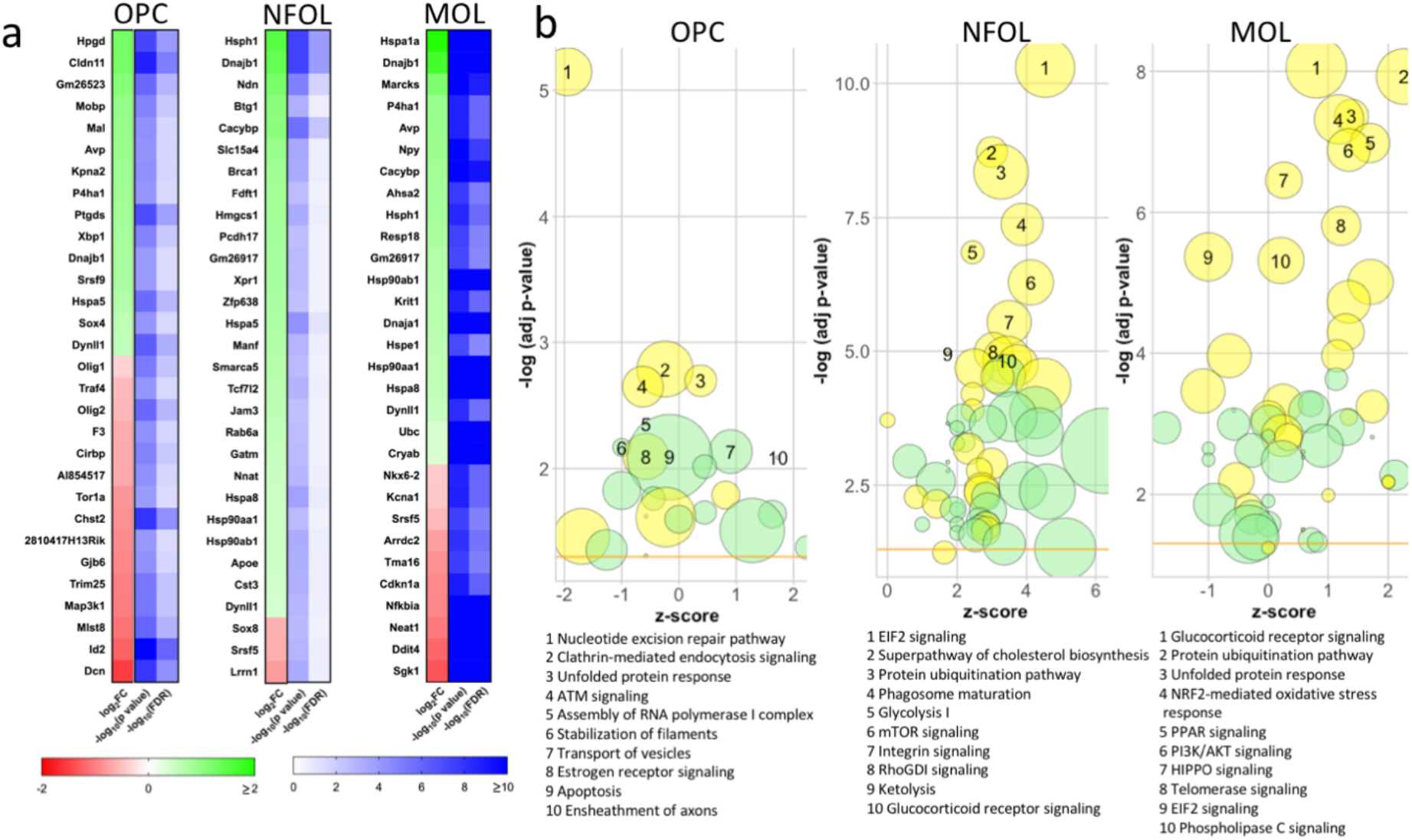
Nutritional signals rapidly regulate the OL transcriptome in the adult ME. (a) Top 30 differentially expressed genes between fasted and refed conditions (*p* < 0.05 and FDR < 0.25, −log_10_(0.05)=1.3) (b) Graphical representation of top 10 pathways (IPA) changed between fasted and refed conditions. Radius indicates number of differentially expressed genes in current dataset that overlap with IPA gene set, yellow = IPA canonical signaling pathway, green = IPA cellular function, yellow horizontal line denotes statistical significance threshold (-log(adj pvalue) of 1.3)

In the OPC cluster, genes implicated in OPC migration (*Cldn11*, *Tspan15*)^13^ were upregulated. Genes regulating OPC proliferation and maintenance in the cell cycle were both activated (*Sox4*, *Hes5*) ^14^ and inhibited (*Id2*, *Gpr17*) ^15^. Likewise, genes promoting OPC differentiation and lineage progression were both activated (*Kpna2*, coordinator of transcriptional control of OL differentiation ^16^) and inhibited (*Olig1, Olig2)* ^15^. These results suggest that both OPC recruitment and proliferation and OPC differentiation may be upregulated in distinct subsets of OPCs in response to the fast-refeed paradigm. Consistent with a transcriptional activation initiating differentiation into OLs, refeeding upregulated transcripts related to peroxisomal activity (*Pex5*, *Pmvk*) and involved in cholesterol and myelin lipid synthesis ^17,18^, and myelin-related transcripts (*Mal, Mobp, Plp1*). Functional annotation of DEGs in OPCs also highlighted genes involved in hormonal and metabolic sensing (*Mlst8, Igf1r, Pik3r1*).

In NFOLs, only 19 genes were significantly differentially expressed between treatments. Strikingly, these include the transcription factor *Tcf7l2,* a key player in OL differentiation albeit with a controversial role on cell cycle exit ^19^. Several transcripts encoding heat-shock proteins that promote survival of myelinating cells were upregulated (*Hsph1*, Dnajb1, *Hspa5, Hspa8, Hsp90ab1*) ^20^, as well as genes involved in lipid/cholesterol synthesis or transport (*Hmgcs1, Apoe*).

Likewise in MOLs, several transcripts for heat shock proteins were consistently upregulated (*Hspa1a, Hsph1, Hspa4l, Hsp90aa1, Hsp90ab1, Hspa8, Hspa5, Cryab, Dnaja1, Dnajb4, Ahsa2*). Three genes mediating cell cycle exit and differentiation (*Klf6*, *Klf9* and *Yy1)*^21 22^ were upregulated; others were downregulated (*Olig1*, *Olig2*, Sgk1 and Cdkn1a) ^23^. Because all these latter genes are enriched in NFOLs compared to OLs ^24^, this could indicate lineage progression to mature OLs. However, the pro-myelinating transcription factor *Nkx6-2*^25^, and the polarity gene *Mtmr2* required for correct myelin formation ^26^were downregulated, while genes promoting lipid biogenesis and cholesterol transport (*Elovl1, Arid5b*, *Stard3*) were upregulated, leaving unclear the consequence of refeeding on OL maturation and myelination.

We also used Ingenuity Pathway Analysis (IPA) to make a comprehensive prediction about pathways regulated by the transition from the fasted to refed state in OL lineage cells of the ME (Figure 4b, Supplementary Table 7). Consistent with the DEG analysis, top pathways included processes involved in proliferation, cell cycle progression, differentiation, lipid/cholesterol biosynthesis and myelination (Supplementary Table 8). In addition, a number of pathways involved in hormonal sensing, growth and cellular energy handling were significantly regulated (Supplementary Table 9). This was particularly pronounced in NFOLs, with a strong activation of EiF2 and mTOR signalling.

Using IPA, we also obtained the top upstream regulators of DEGs, i.e., transcriptional regulators that best explain the expression changes between fasting and refeeding (Supplementary Table 10). Strikingly, top upstream regulators included two transcription factors necessary for OL differentiation, *Tcf7l2* and *Myrf* in all clusters, and *Mtor* in OPCs and NFOLS (Supplementary Figure 4). Corticosterone, glutamate, dopamine and oestradiol were also among the top upstream regulators, introducing a possible regulatory role for these signals in the observed changes.

### Refeeding rapidly triggers the formation of new OLs in the ME

The analysis of DEGs, top differentially regulated pathways, and top upstream regulators indicate that OL lineage progression is regulated during the fast-refeed transition. We used a number of complementary tools to validate these results. With RNAScope FISH, we found that the expression of *Pdgfra* expression decreased while the expression of *Bmp4* and *Tcf7l2* increased in Sox10^+^ cells, resulting in an increase in the number of NFOLs (*Sox10*^+^/*Bmp4*^+^ or *Sox10*^+^/*Tcf7l2*^+^) in response to the refeed (Fig. 5a–5d). The number of MOLs (*Sox10*^+^/*Plp1*^+^) increased in refed samples together with an increase in *Plp1* expression (Fig 5da–5c). Immunodetection of PDGFRa and APC confirmed a decrease in the number of OPCs that was specific to the ventral ME, and an increase in APC^+^ OLs in the ME of refed mice (Supplementary Fig. 5a, Figure 5e,5g). Last, we quantified BrdU incorporation into ME and CC OPCs in fasted and refed mice and examined BrdU colocalization with antibodies to Sox10 and Pdgfrα (Figure 5f). The total number of Brdu^+^/Sox10^+^ cells was not different between conditions in both brain sites (Supp. Fig. 5b). However, in the ME, refeeding produced a decrease in the number of proliferating OPCs (Sox10^+^/Pdgfrα^+^/ BrdU^+^), an increase in the number of newly differentiated NFOLs (Sox10^+^/Pdgfrα^−^/BrdU^+^), and an increase in pre-existing OLs (Sox10^+^/Pdgfrα^−^/BrdU^−^) (Figure 5h). None of these changes were observed in the CC, a white matter tract where rapid OPC differentiation has been characterised before^10,27^ (Figure 5i). In this experiment, we noticed that NFOLs were positioned more dorsal in the ME of refed mice than in fasted controls, suggesting that refeeding also promotes the dorsal migration of OPCs committed to differentiation (Supp. Fig. 5c). Consistent with the idea that activated OPCs ready to be differentiated are migrating dorsally, we found that dorsal OPCs receive significantly less synaptic input than ventral OPCs, a hallmark of OPC differentiation^28^, and that refeeding decreased the number of glutamatergic punta contacting ventral OPCs (Supp. Fig. 5d, 5e). Collectively these data indicate that refeeding after an overnight fast rapidly promotes the formation of new OLs and demonstrate the unique ability of OL lineage cells of the ME to respond to nutritional stimuli.

**Figure 5.**
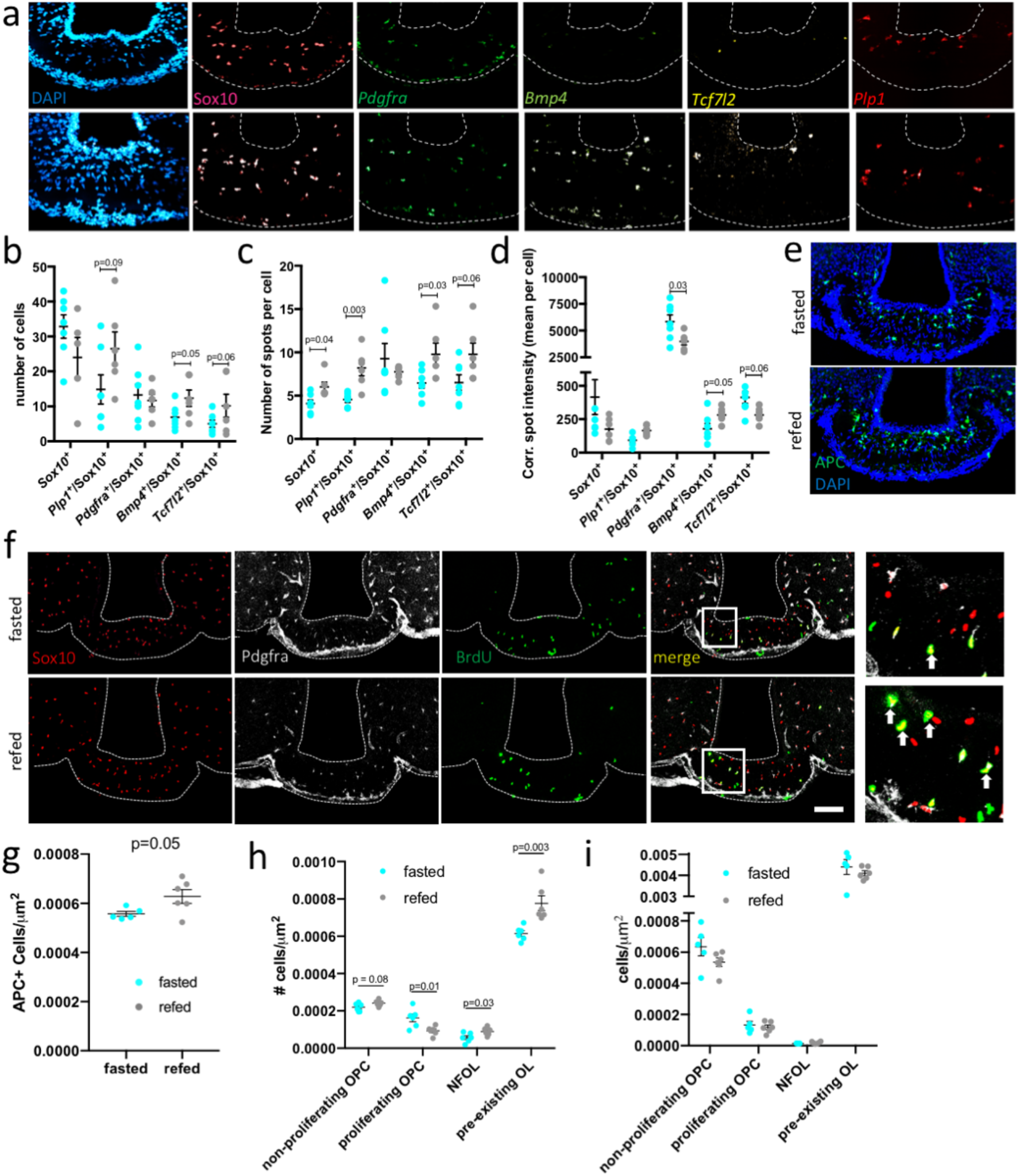
1 hour refeeding increases the formation of new OLs in the ME. (a) Multiplex single molecule FISH labeling OL markers in the ME: red 1 = Sox10 (pan-OL marker), green 1 = *Pdgfra* (OPC marker), green 2 = *Bmp4* (NFOL marker), yellow = *Tcf7l2* (NFOL marker), red 2 = *Plp1* (MOL marker). (b) Number of cells expressing markers, (c) Number of RNA molecules (‘spots’) per cell and (d) corrected spot intensity in FISH experiment. (e) APC (postmitotic OL marker) immunolabelling in ME sections from fasted and refed mice (g) and quantification of the density of APC^+^ cells in this paradigm. (f) Bromodeoxyuridine (BrdU) labelling during in fast/refeed paradigm. Sox10 (red) = pan-OL marker, Pdgfrɑ (grey) = OPCs, and BrdU (green) = cells that have divided during fast refeed paradigm. Squares indicate insets. Arrows denote NFOLs (Sox10^+^/Pdgfr ^−^/BrdU^+^). Quantification of subsets of BrdU labelled cells in the ME h) and the CC (i). Scale bar = 100 μm (d-e) Non-proliferating OPC = Sox10^+^/Pdgfra^+^/BrdU^−^, proliferating OPC = Sox10^+^/Pdgfra^+^/BrdU^+^, NFOL = Sox10^+^/Pdgfra^−^/BrdU^+^, postmitotic OLs before treatment = Sox10^+^/Pdgfra^−^/BrdU^−^. * indicates *p* < 0.05, ** indicates *p* < 0.01, error bars indicate mean ± SEM.

### Acute refeeding does not increase myelination in the ME

Our scRNAseq dataset indicates that many genes translating to proteins involved in myelination are regulated by acute nutritional signals in the ME (Supp. Table 9, Figure 6a, Supplementary Figure 6). Expression of genes involved in cholesterol, phospholipid, triglyceride and sphingomyelin is increased in the NFOLs population of the ME compared to OPCs and MOLS (Supplementary Figure 6). Collectively these results suggest that refeeding could produce a rapid increase in myelination in the ME.

**Figure 6.**
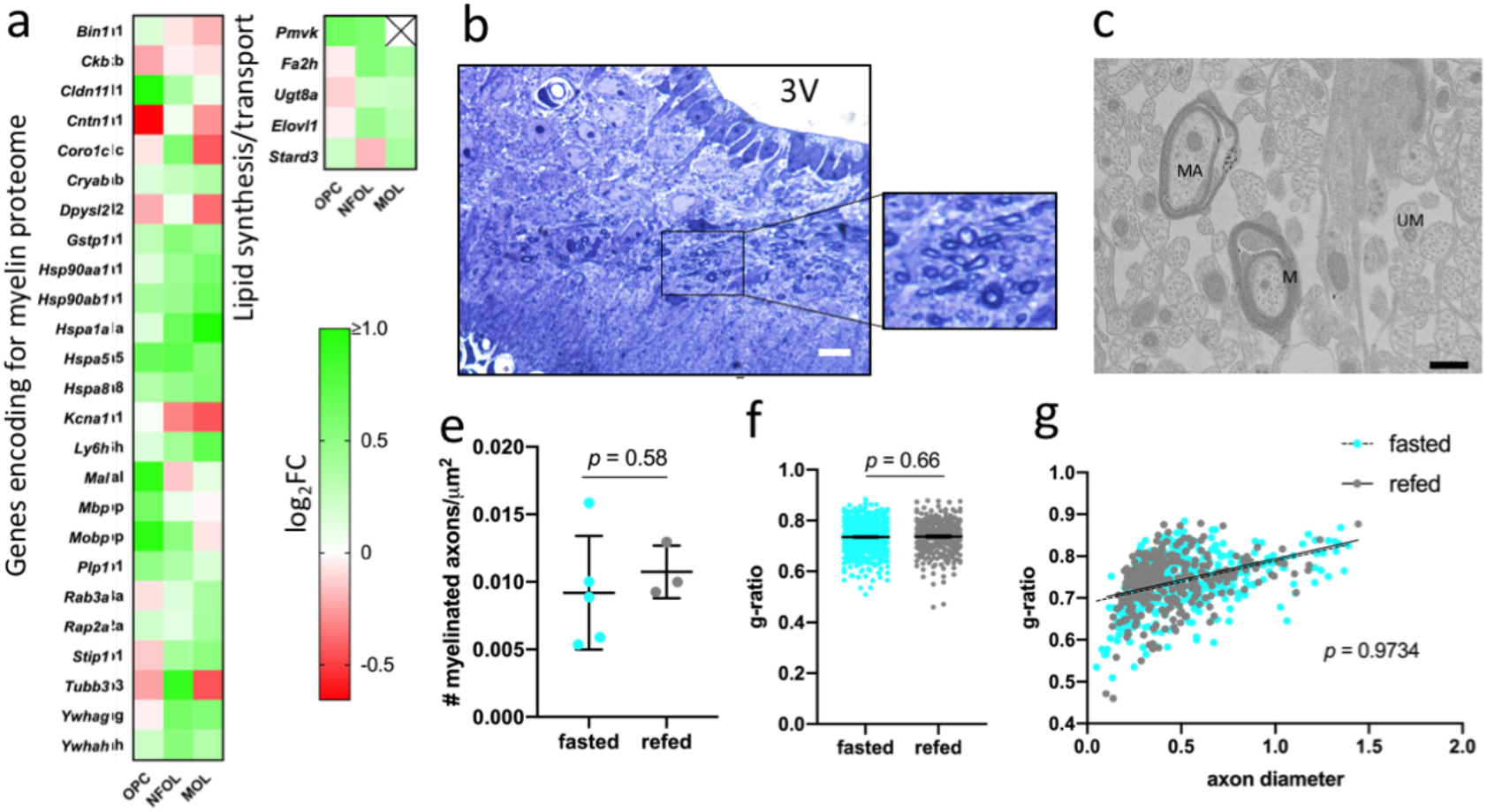
Myelination does not increase after a 2 hour refeed. (a) Log_2_FC of DEGs (*p* < 0.05 and FDR < 0.25) translating to proteins of the myelin proteome or involved in lipid synthesis and transport. An ‘X’ indicates the gene is not expressed in one or both conditions in that OL cluster. (b) Toluidine blue-labelled of thin ME sections. 3V = third ventricle, scale bar = 10 μm, box indicates inset. Dark circles are transverse cross sections of myelin ensheathing an axon. (d) Transmission electron micrograph of myelin in the ME. Scale bar = 500 nm, M = myelin, MA = myelinated axon, UM = unmyelinated axon. (e) Measurements of density of myelinated axons in the ME of animals fasted (n = 5) or refed (n = 3) mice (f) Measurements of g-ratio of myelinated axons in fasted (n = 5 mice, 481 axons) and refed (n = 3 mice, 308 axons) conditions. Error bars indicate mean ± SEM (g) Measurement of slopes of linear regression lines to fit g-ratio (as in f) plotted against axon diameter

To determine if the increase in the expression of genes related to lipid biosynthesis and myelin transcription translates to changes in myelination in this acute paradigm, we fasted animals overnight and refed them for 2 hours and used Toluidine blue labelling of thin sections to view the distribution of myelinated axons in the ME. In line with MBP immunolabelling in thick cleared sections (Video 1), myelinated axons are seen in the transverse plane, indicating their passage in a rostral-to-caudal direction through the ME (Figure 6b-c). In addition, myelinated axons are all localised to the dorsal third of the ME where MBP and *Plp1* are expressed (Figure 6b, Figure 3a, d). Using transmission electron microscopy, we quantified the density of myelinated axons and myelin thickness. None of these changed during a 2h refeed following an overnight fast (Figure 6b-g), suggesting that in this time frame, changes in myelination do not occur.

### OPC and OL response to a fast-refeed modifies the extracellular matrix of the ME

The lack of measurable changes in myelin-related parameters in the ME of refed mice left unaddressed the functional relevance of the rapid nutritional regulation of OLs in the ME. This prompted us to consider alternative mechanisms through which changes in OL differentiation during the fast-refeed transition may regulate the function of the ME. We focused on DEGs encoding secreted proteins to identify potential nutritionally-regulated paracrine pathways through which OPCs and OLs may communicate with other functionally-relevant cells types of the ME or modify the local ME environment. This revealed that a number of genes expressing proteins of the extracellular matrix (ECM) were among top DEGs, including several proteoglycans. In OPCs, the gene most significantly regulated by refeeding is *Dcn* (log_2_FC_fast→refeed_=−1.5, p-value=1.6E-08, FDR=4.22E-05), encoding decorin, a proteoglycan involved in decreased barrier properties of the blood-brain barrier in response to brain injury^29^. Interestingly, we found that OL subtypes are sole or main producers of other ECM proteins including key components of perineuronal nets (PNNs) tenascin-R (TNR) and veriscan (Vcan) (both highly enriched in NFOLs, Fig. 7a and 7b). PNNs have been recently shown to enmesh key metabolic sensing neurons at the ARC-ME junction, and proposed to regulate how these neurons access blood-borne signals^30^. To determine whether the increased number of NFOLs in refed mice would change the expression of TNR or the density of PNNs, we performed immunolabelling against TNR and Wisteria floribunda agglutinin (WFA), a lectin specifically labelling PNNs, and quantified volumes and number of labelled objects using high resolution confocal imaging and 3D reconstruction of 20uM-thick stacks. In line with the increased number of NFOLs, the main source of TNR, we found that the total volume of TNR labelling and the number of TNR-labelled surfaces was significantly higher in the ME of refed mice (Fig. 7c and 7d). This was accompanied by a significant increase in the number of WFA-labelled surfaces in the ME of refed mice, supporting an acute nutritional regulation of PNNs in this paradigm (Fig. e and 7f). Thus, nutritional regulation of OLs produces changes in the local ME extracellular matrix which could change how local metabolic sensing neurons access blood-borne signals.

**Figure 7:**
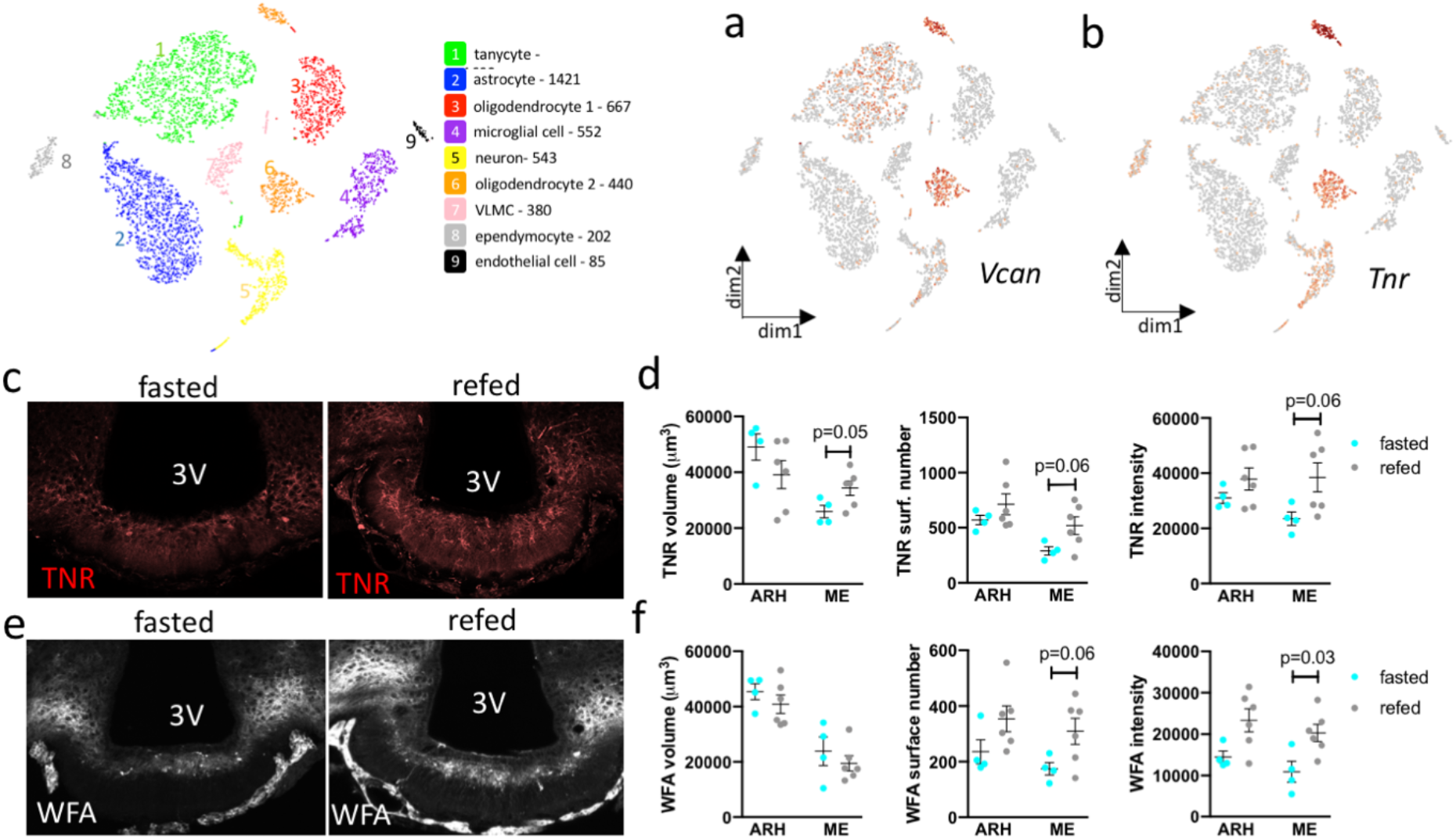
OPC and OL response to a fast-refeed modifies the extracellular matrix of the ME and perineuronal nets. (a) tSNE plots of ME scRNAseq dataset showing expression Vcan and TNR and their enrichment in cluster 6 (oligodendrocyte cluster 2 containing OPCs and NFOLs). Red = high expression, grey = low expression. (c, e) Maximum projection stacks of 20uM ME sections immunolabelled against TNR and WFA and (d, f) volumetric, surface number and intensity quantification in these stacks. Data are means ± sem.

### mTORC1 is highly active specifically in OLs in the ME, is rapidly upregulated during the fast-refeed transition, and responds to dietary protein intake

The IPA results highlighting a role for mTOR in the rapid nutritional regulation occurring in ME OLs prompted us to further investigate the regulation of this pathway in the fast-refeed paradigm. Consistent with IPA, several genes involved in the mTOR signalling pathway (*Igf1r, Pik3r1, Mlst8* which negatively regulates mTOR activity*, Sgk1, Ddit4,* and *Yy1)* are significantly regulated by the refeeding stimulus (Figure 8a, Supplementary Fig 7a). mTOR signalling plays a key role in OL differentiation, maturation and in myelination^31^. Consistently, mTOR was one of the top pathways regulated in NFOLs by the fast-refeeding transition (Figure 4b, Supp. Table 9) – a cell type that actively prepares to make myelin. mTOR signalling is also a key cellular integrator of nutritional and hormonal signals in the regulation of many anabolic processes ^32^, raising the possibility that mTOR signalling in OLs couples metabolic sensing to changing in OL differentiation. Of note, OPCs, NFOLs and OLs express genes to receptors for many of the metabolic hormones that signal energy availability to the hypothalamus, including Adipor2, Fgfr2, Inr, Lepr, Ghr, Gipr and Thra (Supp. Fig. 7b), many of which have been shown to activate hypothalamic mTOR signalling^33–36^.

**Figure 8.**
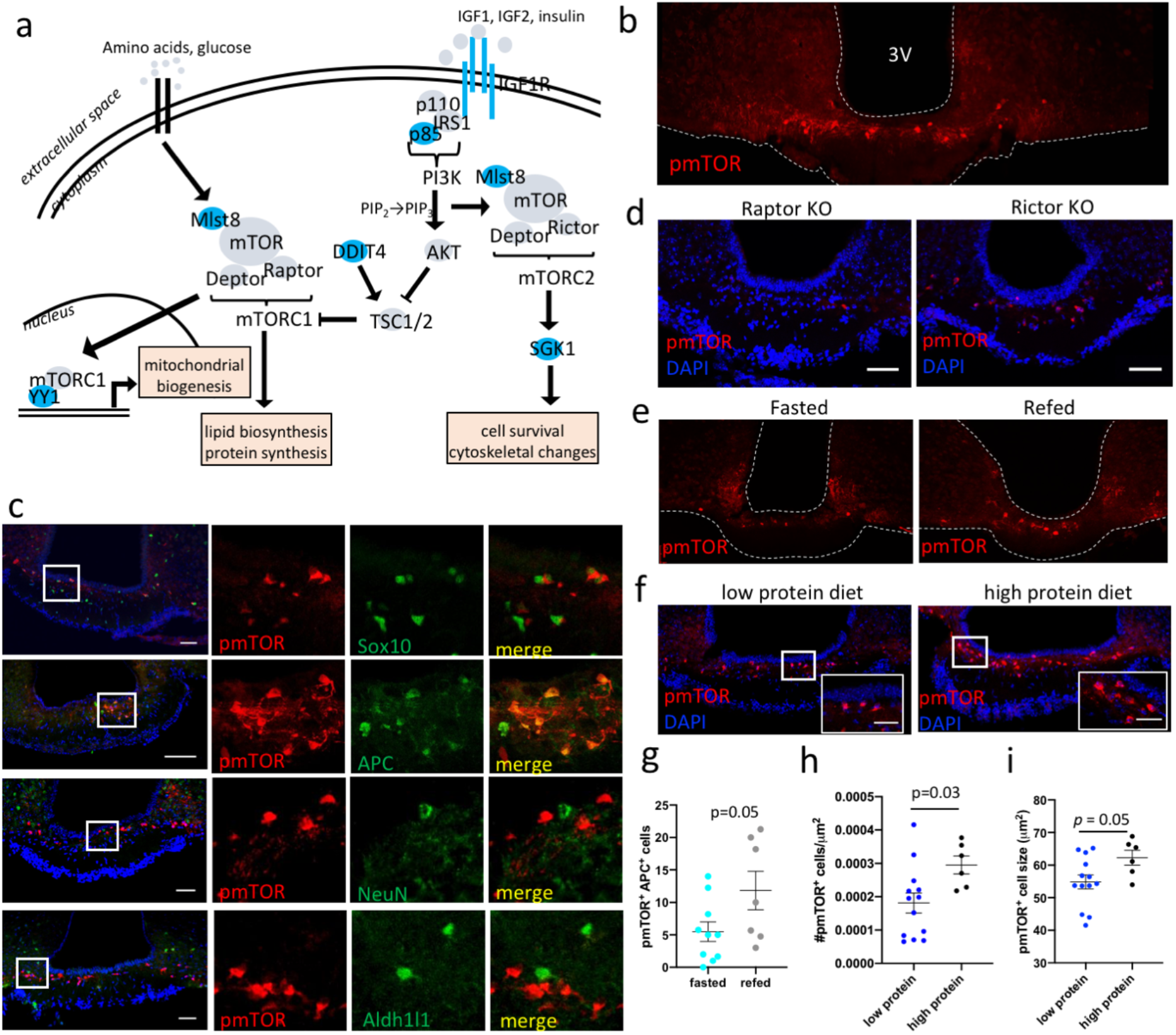
mTORC1 signaling is highly active and nutritionally regulated in the ME OLs. (a) Schematic of mTOR signaling pathway with DEGs labelled in blue (b) Immunolabelling for phosphorylated mTOR (pmTOR) is highly enriched in the ME (c) Cells labelled with an antibody to pmTOR (red) colocalize with cells labelled by pan-OL marker Sox10 and postmitotic OL marker APC (green), but not with cells labelled by neuron marker NeuN or astrocyte marker Aldh1l1 (green) Scale bars = 100μm, white squares indicate insets (d) mTOR labelling in tissue with absence of mTORC1 signaling (raptor KO) or absence of mTORC2 signaling (rictor KO). Scale bars = 100μm. (e, g) pmTOR immunolabelling in the ME of mice fasted overnight or refed for 1h. (f, h, i) pmTOR immunolabelling in the ME of mice fed a low or high protein diet. White squares indicate insets, scale bars = 25μm. Error bars indicate mean ± SEM.

We first assessed the expression of the active form of mTOR in the mouse ME using an antibody detecting the mTOR phosphorylated at Ser2448 (pmTOR). We discovered that pmTOR immunolabelling in the ME was remarkably bright in a cell population lying in the dorsal ME and higher than anywhere else in the Hypothalamus (Figure 8b). Similar labelling of pmTOR was absent from the rest of the brain, including other circumventricular organs (Supplementary Figure 7c). To confirm the identity of the cells expressing pmTOR, we colocalised pmTOR with established markers of OLs, neurons, and astrocytes. ME pmTOR immunolabelling was absent from neurons and astrocytes but colocalised with the pan-OL marker Sox10 (Figure 8c). 37.3 ± 4.54% of Sox10^+^ cells were pmTOR^+^, showing that only a subset of OLs of the ME have active mTOR signalling. In fact, the ME pmTOR^+^ signal highly colocalized with the postmitotic OL marker APC (Figure 8c) indicating that NFOLs or MOLs and not OPCs have active mTOR signalling.

mTOR can signal via two different complexes, resulting in the activation of distinct transcriptional programmes including protein/lipid synthesis or cytoskeletal organization^21^. To investigate if mTOR signalling was occurring via the mTOR Complex 1 (mTORC1, contains raptor protein) or Complex 2 (mTORC2, contains rictor protein), we took advantage of exclusive expression of 2’,3’-cyclic-nucleotide 3’-phosphodiesterase (CNPase) in OLs and used CNP-Cre:raptor^fl/fl^ (Raptor knockout) and CNP-Cre:rictor^fl/fl^ (Rictor KO) mice to interrogate which signalling complex is most active in OLs of the ME. When we labelled tissues of raptor or rictor KO mice with the antibody to pmTOR, we were only able to detect an absence of labelling in raptor KO mice (Figure 8d) thus indicating that mTORC1 signalling is highly active in ME OLs.

As our scRNAseq dataset shows mTOR signalling is regulated by fasting and refeeding in ME OLs, we tested whether mTOR activity in ME OLs would change in this paradigm. We found that refeeding significantly increases the number of pmTOR^+^ OLs (Figure 8e, 8g). In addition, there was a trend for an increase in cell size in pmTOR^+^ OLs (p=0.08) and increased signal intensity (p=0.07).

Many nutritional and hormonal signals in the refed state could potentially produce increased mTORC1 activity in OLs. To further test the idea that nutritional signals are sensed via mTORC1 in OLs to promote OL differentiation, we assessed the effect dietary proteins on OL differentiation. This choice was based on the fact that dietary protein is one of the strongest nutritional signals that activates mTORC1 signalling in multiple tissues. In addition, *Bckdk*, which encodes one of the key enzymes in branched-chain amino acid metabolism, is one of the TOP upstream regulator explaining the transcriptional changes during the fast refeed transition in MOLs (Supplementary Fig. 8). Mice were exposed to a low or high protein diet for 4 days, a time frame during which these diets do not differentially impact weight gain (not shown). We found that high protein feeding increased the density of pmTOR^+^ OLs and increased the size of pmTOR^+^ cells (Figure 8f, 8h-i). Thus, amino acid availability regulates mTORC1 signalling in OLs.

## Discussion

Our study provides a high-resolution transcriptional characterisation of distinct cell types of the ME and their response to nutritional transitions, allowing the identification of cell type-specific cellular processes regulated by nutrient availability. We focused our analysis on oligodendrocytes of the ME, a cell type previously uncharacterised in this brain region. We can conclude from our experiments that ME OLs are structurally highly organised and plastic in adulthood. This plasticity is sensitive to nutritional signals, at least in part via the activation of the mTORC1 pathway in OLs, leading to an increase in OL differentiation. We show that dietary protein is sufficient to activate OL mTORC1 signalling, and that the increased production of adult-born OLs in response to nutrients is associated with changes in extracellular matrix components and increased density of PNNs in the ME-ARC barrier. Collectively, these results identify a novel mechanism through which nutrients regulate ME function and reveal the high sensitivity of OPC differentiation to unbuffered blood-borne nutritional signals.

Our data support a role for mTORC1 in OLs as a nutrient sensor surveying extracellular nutrient availability and coupling OL differentiation to substrate availability. Alternatively, changes in local neuronal activity in response to refeeding could indirectly increase mTORC1 activity and OL differentiation{Saxton, 2017 #609}. The regulation of OL mTORC1 by dietary protein content indicates that mTORC1 in ME OLs directly responds to nutrients. This supports the interpretation that the uniquely high mTORC1 expression in OLs of the ME and unique nutritional regulation of ME OLs, at least compared to OLs of the CC, are a consequence of the exclusive environment of the ME with unbuffered access to peripheral nutrients and metabolic signals.

The high density of NFOLs in the human and murine ME raises the question of the role of OL plasticity in ME functions. In the rest of the brain, OLs are long-lived and adult-born OLs add myelin to existing structures{Hughes, 2018 #398}{Tripathi, 2017 #396}. Further studies are needed to characterise the stability of ME OLs and ME myelin, and determine whether NFOLs in the ME survive long-term, replace mature OLs that turnover with time, or represent a transient population with myelin-unrelated roles. These could include the regulation of the ME extracellular matrix and PNNs, as suggested by our results. More experiments will be needed to directly demonstrate the role of NFOLs in the ME-ARC barrier and how this may alter the activity of ARC neurons. In spite of the lack of measurable changes in myelin-related parameters in response to acute refeeding, the very high mTORC1 activity in ME OLs after refeeding, the increase in OL cell size, and the observed transcriptional changes indicate that these cells are actively preparing for myelination. More chronic nutritional manipulations may be required to see these changes.

Our results provide novel knowledge on the regulation of OLs by nutritional signals, raising the question of how OLs of other brain sites respond to blood-borne signals when the BBB is compromised, such as after brain injury. Interestingly, in this pathological context, both increased NFOL production and remodelling of the extracellular matrix are occurring^37^. Whether remyelination post-injury is regulated by peripheral energy availability has not been determined to our knowledge. Coupling OL differentiation to energy availability seems appropriate, given the high energetic cost of myelination and requirement for specific nutrients. Extensions of our findings to these models may provide novel nutrition-based strategies to optimise remyelination after injury and better understand the regulatory role of the nutritional and metabolic status on OL biology.

## Supporting information

Merged pdf od supplementary figures

All supplementary tables

## Acknowledgments

We thank Chiara Cossetti at the Flow Cytometry Core of the Cambridge Institute of Medical Research for assistance with FACS. We thank James Warner at the Wellcome Trust-MRC Institute of Metabolic Science Histopathology core for his assistance with processing human tissue. We thank Gregory Strachan at the Wellcome Trust-MRC Institute of Metabolic Science for his assistance with confocal imaging. We thank Fadwa Joud at the Light Microscopy Core at the Cancer Research UK Cambridge Institute for her assistance with microscopy. We would like to thank the Cambridge Brain Bank (CBB) for providing brain tissue for this study. This work was supported by the Medical Research Council [MR/S011552/1](CB), the Wellcome Trust four-year PhD Programme in Metabolic and Cardiovascular Disease [108926/B/15/Z](SB), the BBSRC DTP programme (SK), the Medical Research Council Metabolic Disease Unit and the Wellcome Trust Strategic award for the MRL Disease Model Core and Imaging facilities [MRC_MC_UU_12012/5], [100574/Z/12/Z], [MRC_MC_UU_00014/5], [208363/Z/17/Z], the Adelson Medical Research Foundation (RJMF and DHR), the UK MS Society (RJMF and CZ) and the l’Oreal-UNESCO For Women In Science programme.

## Author contributions

SK, BL, SB, CZ, DN, JT, SH, KR, HH and CB performed experiments. SK, BL, SB, CZ, DN, JT, SH, KR, HH and CB performed data analysis. BL, WM, GSHY and DR designed experiments. SK, RJMF, DR and CB wrote the manuscript.

## Methods

### Animals

7-9-week-old male C57BL/6 mice were used in all studies except where otherwise noted. Mice were maintained on a 12h light/dark cycle, had free access to water and normal chow (Safe Diets – Safe 105m) and were group-housed (at least 2 animals per cage). Animals were handled regularly before experiments to reduce stress-related responses. All studies were approved by the local Ethics Committee and animals were treated in accordance with the UK Home Office (Scientific Procedures) Act (1986). CNP-Cre:raptor^fl/fl^ and CNP-Cre:rictor^fl/fl^ brain tissue was provided by Professor Macklin from the University of Colorado Anschutz Medical Campus. Aldh1l1-GFP mice were provided by Professor Tchöp from the Helmholtz Diabetes Center & German Center for Diabetes Research.

### Fast-refeed paradigm

Food was removed just before dark onset and for 16h. Access to food was then restored in refed animals for up to 2h before sacrifice. For the BrdU labelling experiment only, animals began fasting at noon and refed animals were fed at 11am the next day.

### Low protein/high protein paradigm

After 3 consecutive brief exposures to the novel diets to avoid neophobia, mice were fed isocaloric diets containing either 7% or 45% of energy as casein for 4 days (Research diets, LP: D17030701, HP: D17030703, respectively). The LP diet consisted of 20% kcal fat (from soybean oil), 73% kcal carbohydrate (from corn starch and sucrose), and 7% kcal protein (from casein). The HP diet consisted of 20% kcal fat (from soybean oil), 35% kcal carbohydrate (from corn starch and sucrose), and 45% kcal protein (from casein).

### Bromodeoxyuridine (BrdU) administration for fast-refeed experiment

Mice were fasted at noon and received 2 ip injections of BrdU (Sigma, 50 mg/kg in saline), at 1pm and 5pm respectively. The next day they received 2 additional BrdU injections at 9am and 11am, followed by a refeed of 1h or an additional 1h of fast, and sacrifice at 1pm.

### Perfusion fixation

Animals were anaesthetized with an ip injection of 50 ul pentobarbitol (Dolethal, 200 mg/ml) then transcardially perfused as follows. For immunohistochemistry (IHC), tissue clearing, and RNAscope experiments, animals were perfused with 0.01 M phosphate buffered saline (PBS) at room temperature (RT) followed by 4% paraformaldehyde (PFA, Fisher Scientific) in PBS (pH 7.4) at 4°C. For experiments requiring resin embedding, animals were perfused with cold 4% glutaraldehyde (Generon), 0.008% CaCl2 (Sigma) in PBS.

## Single-cell RNA sequencing

All data are available on GEO (GSE133890).

### Tissue dissection and dissociation

Tissue dissociation for single-cell RNA sequencing (scRNAseq) was performed as previously described^38^. 10 P40-47 mice were fasted overnight, half were refed for 1h as described above. Animals were sacrificed via cervical dislocation and the brain was quickly extracted into cold Neurobasal-A medium (ThermoFisher Scientific). The ME was dissected from each brain with fine curved scissors (Fine Science Tools - No. 15010-11). MEs were placed in ice cold papain (Worthington Biochemicals - LK003160, 20 U/ml in Hibernate A) in separate 1.5 ml Eppendorfs until all dissections were finished. The sections in papain were incubated at 37°C (500 rpm) for 15-20 min, and the Eppendorfs were swirled every 5 min. The tissue was then triturated in prewarmed DNase solution (Sigma-Aldrich - D4263) at 37°C then placed in tubes on ice until fluorescence-activated cell sorting (FACS).

### Fluorescence-activated cell sorting

The cell suspensions from triturations were passed through a 40 um cell strainer into fresh collection tubes. DraQ5 and DAPI were added to the samples in DNase solution to select for nuclei and exclude dead cells, respectively, then single cells were sorted with an Influx Cell Sorter (BD Biosciences) into tubes containing 10 ul 0.4% BSA in Ca-/Mg-free PBS. 3500 cells were sorted into each tube then kept on ice until sequencing.

### Sequencing

Isolated cells were encapsulated in droplets and cDNA libraries were made using a 10X Genomics Chromium instrument and 10X Single Cell 3’ V2 Reagent kit. Paired end sequencing was performed on an Illumina HiSeq 4000. The first 26bp read contains both a cell barcode and a unique molecular identifier, the next 76bp read contains the cDNA insert. The sequencing reads were mapped to the Genome Reference Consortium m38 (mm10) mouse reference genome and counted using 10X Genomics Cell Ranger software version 2.0. Library preparation and sequencing was performed at the Genomics Core, Cancer Research UK Cambridge Institute. Sequencing data are available on GEO (accession number GSE133890). Of 14,000 cells captured via FACS, 5,982 cells were successfully sequenced using the 10X platform Single Cell 3’ v2. A median of 1,853 genes were expressed per cell, with a median of 3,837 UMI counts per cell.

## Bioinformatics

All code used for R analysis of scRNAseq data is found in the online repository Github: Kohnke-et-al-2019.

### T-distributed stochastic neighbor embedding and clustering

R software was used to analyse scRNAseq data. The R package ‘cellrangerRkit’ (supported by 10X Genomics) was used to perform t-distributed stochastic neighbor embedding (TSNE), a dimensionality reduction technique that allows mapping of all cells in 2 dimensions based on their transcriptomic profile. The package ‘NBClust’ ^38^ was used to test the TSNE plot for the optimal number of clusters. Finally, cellrangerRrkit was used to find the top defining genes per cluster.

### Differential gene expression

The R package ‘edgeR’^39^ was used to identify differentially expressed genes (DEGs) between fasted and refed conditions in individual clusters. This package fits gene expression data from one cluster from one condition to a generalized linear model (GLM) then compares it to expression in the same cluster in the other condition. GLM fitting is beneficial for complex multifactor experiments. edgeR generated an output of log2-fold change (log2FC) in expression, p-values, and false discovery rates (FDRs) for every gene of each cluster.

### Pathway analysis

Ingenuity Pathway Analysis (Qiagen) was used to identify pathways that are up- or downregulated in each cluster between experimental conditions. Log2FC values, p-values, and FDRs for each gene for each cluster were uploaded to the software. Results were confirmed with DAVID functional annotation (data not shown) ^40,41^.

## Fluorescence in situ RNA hybridization

### Mouse tissue

Brains were postfixed in 4% PFA solution overnight then cryoprotected in 30% sucrose solution in PBS for up to 24h. Tissue was covered with optimal cutting temperature (OCT) media then sliced at 16 μm thickness using a Leica CM1950 cryostat directly onto Superfrost Plus slides (ThermoScientific) in an RNase free environment. Slides were then stored at −80°C. Sections were sliced in the coronal plane from Bregma −1.58 to −2.30 mm^42^.

Fluorescence multiplex in situ RNA hybridisation (FISH) was performed as previously described using RNAscope technology{Bayraktar, 2018 #359}^43^. After epitope retrieval and dehydration, sections on slides were processed for multiplexed FISH using the RNAScope LS Multiplex Assay (Advanced Cell Diagnostics) followed by immunohistochemistry on a Bond RX robotic stainer (Leica). Samples were first permeabilised with heat in Bond Epitope Retrieval solution 2 (pH 9.0, Leica - AR9640) at 95°C for 2 min, incubated in protease reagent (Advanced Cell Diagnostics) at 42°C for 10 min, and finally treated with hydrogen peroxide for 10 min to inactivate endogenous peroxidases and the protease reagent. Samples were then incubated in z-probe mixtures (*Pdgfra* 1:1, *Bmp4* 1:50, *Tcf7l2* 1:50 and *Plp1* 1:400) for 2 h at 42°C and washed 3 times. DNA amplification trees were built through incubations in AMP1 (preamplifier), AMP2 (background reducer), then AMP3 (amplifier) reagents (Leica) for 15-30 min each at 42°C. Between incubations, slides were washed with LS Rinse buffer (Leica). After, samples were incubated in channel-specific horseradish peroxidase (HRP) reagents for 15 min at 42°C, tyramide signal amplification (TSA) biotin or TSA fluorophores for 30 min and HRP blocking reagent for 15 min at 42°C. The following TSA labels were used to visualize z-probes: Atto 425-streptavidin (Sigma - 40709, 1:200), Opal 520 (1:500), Opal 570 (1:500), and Opal 650 (1:2500) fluorophores (Perkin Elmer).

Directly following the FISH assay, tissue was incubated with anti-Sox10 antibody in blocking solution for 1 hour (Abcam - AF2864, 1:100). To develop the antibody signal, samples were incubated in donkey anti-goat HRP (Thermo fisher Scientific - A15999, 1:200) for 1 hour, TSA biotin (Perkin Elmer - NEL700A001KT, 1:200) for 10 min and streptavidin-conjugated Alexa 700-streptavidin (Sigma - S21383, 1:200) for 30 min.

### Human tissue

Human hypothalamic tissues used for this study were from donors to the Cambridge Brain Bank. Donors gave informed written consent for the use of brain tissue for research and tissues obtained were used in accordance with the Research Ethics Committee Approval number 10/H0308/56. Samples were from an 83-year-old with no neuropathology. 59h after death, hypothalamic samples were collected and stored in 10% neutral buffered formalin at room temperature for 24h, transferred to 70% ethanol, and processed into paraffin. 6 μm sections were cut and mounted onto Superfrost Plus slides (Thermo-Fisher Scientific) in an RNase free environment, and then dried overnight at 37°C.

Using a Bond RX robotic stainer (Leica), slides were deparaffinized, rehydrated, treated with Epitope Retrieval solution 2 88°C for 15 min, and with ACD Enzyme from the Multiplex Reagent kit at 40°C for 10 min. Z-probes (PDGFRA 1:50, ACD - 604488-C4; BCAS1, 1:50, ACD - 525788-C3; PLP1, 1:1, ACD - 499278) were used to detect mRNA transcripts in the tissue. Probe hybridisation and signal amplification was performed according to manufacturer’s instructions. The following TSA labels were used to visualize z-probes: TSA plus-Cy5 (1:750 to detect PLP1, Akoya Biosciences - NEL745001KT), TSA plus-Fluorescein (1:300 to detect BCAS1, Akoya Biosciences - NEL741001KT) and Opal 620 (1:300 to detect PDGFRA, Akoya Biosciences - FP1495001KT). After completion of the FISH assay, slides were removed from the Bond RX and mounted using Prolong Diamond (ThermoFisher - P36965). We thank Julia Jones at the histopathology/ISH core facility at Cancer Research UK-Cambridge Institute for assistance with in situ hybridisation.

### Immunofluorescence

Brains were postfixed in 4% PFA solution overnight then cryoprotected in 30% sucrose solution in PBS for up to 24h. Tissue was covered with optimal cutting temperature (OCT) media (CellPath), and 30 μm thick coronal sections were obtained from Bregma 0.62 to −7.76 mm^39^ using a Bright Series 8000 sledge microtome.

Antigen retrieval was used for all experiments prior to antibody incubation. Sections were incubated in 10 mM sodium citrate (Fisher Scientific) in distilled water at 80°C for 20min then washed 3 times in PBS. For stains including the BrdU antibody, sections were then incubated in 2 N hydrochloric acid (Sigma) in distilled water at 37°C for 30min. The acid was neutralized by washing sections in 0.1 M sodium tetraborate (Sigma) in distilled water with hydrochloric acid to adjust pH (final pH = 8.5) for 10min, then sections were washed 3 times in PBS. For all experiments, sections were blocked in normal donkey serum (NDS, Vector Biolabs) in PBS plus 0.3% Triton X-100 (Sigma-Aldrich, 0.3% PBST) for 1h prior to primary antibody incubation (antibodies were diluted in 0.3% PBST with or without block).

Sections were incubated in primary antibody solution (Supplementary Table 1) for the appropriate time at 4°C, then washed and incubated with appropriate secondary antibodies (Supplementary Table 2) diluted at 1:500 in 0.3% PBST for 90min at RT. Sections were then washed and mounted on slides (Clarity) with mounting media containing 4ʹ,6-diamidino-2-phenylindole (DAPI, Life Technologies Corporation) and covered with thickness 1.0 coverslips (Marienfeld).

## Tissue clearing

Brains were postfixed in 4% PFA solution overnight. Whole brains were washed in PBS 2 times for 2h. Brains were trimmed in the coronal plane using a Leica VT1000s vibrotome until approximately Bregma −1.58 mm. Four to six 200 μm sections were then sliced from each brain and placed in a 24- well plate (Costar) in PBS. Tissue clearing was performed using the Clear, Unobstructed Brain/Body Imaging Cocktails (CUBIC) method as published^44,45^, with minor modifications.

### Preparing reagents

The CUBIC1 solution containing 25% urea (VWR Chemicals – 443874G), 28.8% distilled water, 31.2% Quadrol (Aldrich - 122262, diluted to 80% in distilled water) and 15% Triton X-100 (Fisher Bioreagents – BP151) was made by mixing the first 3 components on a hot plate at 150°C for 15min. After cooling, Triton-X 100 was added to the solution and mixed at RT. CUBIC2 solution is comprised of 25% urea, 50% saccharose (VWR Chemicals – 443815S), 15% distilled water, and 10% triethanolamine (Sigma – 90279). Urea, saccharose, and water were mixed on a hot plate at 150°C for 30min then allowed to cool to RT. Triethanolamine was then added to the solution and mixed at RT.

### Tissue clearing – CUBIC1

PBS in the well plate was replaced with a mixture of 1:1 CUBIC1 solution and distilled water plus Hoechst stain (Life Technologies – H3570, 1:2000, 1ml per well). The seam of the well plate was sealed with Parafilm and the well plate was placed in a shaking waterbath at 37°C for 3h. The CUBIC1/water solution was then discarded and replaced with 100% CUBIC1 solution plus Hoechst and kept at 37°C overnight. The following day, the samples were washed in PBS three times for 1h on a shaker. The samples were then placed in 30% sucrose in PBS until the sections sank to the bottom of the wells. After, the sections were immersed in OCT and frozen at −80°C at least overnight.

### IHC labelling of cleared tissue

The samples were thawed and washed in PBS three times for 1h on a shaker. The samples were placed in a new 24-well plate and covered with primary antibodies (Supplementary Table 1) diluted in PBS plus 2% Triton X-100 (2% PBST) and 10% NDS for 48h at 4°C on a shaker. Samples were then washed in 0.3% PBST at RT three times for 1h. The samples were then placed in secondary antibodies (Supplementary Table 2) diluted in 2% PBST and 10% NDS for 48h at 4°C on a shaker. Samples were then washed in 0.3% PBST at RT three times for 1h.

### Matching refractive index – CUBIC2

IHC-labelled samples were immersed in a mixture of 1:1 CUBIC2 solution and PBS. The well plate was sealed with Parafilm then placed in a shaking waterbath at 37°C for 3h. The CUBIC2/water solution was then discarded and replaced with 100% CUBIC2 solution and kept at 37°C at least overnight, but maximum 72h. The day before imaging, samples were placed in a 1:1 mixture of mineral oil (Sigma-M8410) and silicone oil (Sigma - 175633).

## Myelin analysis

### Resin embedding

Brains were postfixed in 4% glutaraldehyde for 24h then moved to PBS. 1 mm-thick sections containing the ME were sliced by hand from the brains and were stained with 2% osmium tetroxide (Oxkem) for 24h at 4°C. Sections were washed with water 3 times then dehydrated using an ethanol gradient as follows, on a rotator: 50% 2 times for 15min, 70% 2 times for 15min, 90% 2 times for 15min, 95% 2 times for 15min, 100% 3 times for 10min. Sections were then placed in propylene oxide (Agar Scientific) for 20 min. Sections were incubated in a mixture of 1:1 propylene oxide and resin (TAAB) for 6h on a rotator, then in 100% resin for 24h. Sections were mounted in resin in plastic molds (TAAB) and incubated at 60°C for 24h.

### Toluidine blue labelling

Resin blocks were trimmed with a microtome (Leica RM-2065) to expose the tissue then 0.75 μm-thick sections were placed on a water droplet on a slide. Slides were heated on a hotplate to evaporate the water, then toluidine blue (0.5%, Merck) was applied to the sections for 30 seconds before washing off with distilled water.

### Post staining for transmission electron microscopy

Resin embedded tissues were trimmed around the ME and semi thick slices were cut to create sections that only contained the ME. Then, 70nm ultrathin sections were sliced on an ultramicrotome (Reichert-Jung - Ultra-cut 701701 Ultra Microtome) with a diamond knife (Diatome-Ultra 45). Sections were placed on mesh copper grids (size 300) and were post stained with aqueous uranyl acetate for 6 min then lead citrate for 2 min.

## Imaging

### Confocal microscopy

Thin immunolabelled mouse sections (63x and 40x oil objective) and human tissue labelled with FISH (10x dry and 20x oil objectives) were imaged using a Leica SP8 confocal microscope. For mouse tissue, sections were imaged at multiple points in the z plane (z-stacks) at intervals of 3.3 μm to collect signal from the entire depth of the tissue for the region of interest (ROI). ROIs for mouse tissue included the vascular organ of the lamina terminalis, subfornical organ, median eminence (ME), arcuate nucleus of the hypothalamus (ARC), corpus callosum (CC), and area postrema. For human tissue, sections were imaged at multiple points in the z plane (z-stacks) at intervals of 5 μm or 0.5 μm for volumetric analysis. ROIs for human tissue were the ME and ARC at 20x and the full coronal hypothalamic section at 10x. Gain and laser power settings remained the same between experimental and control conditions within each experiment.

### High-content confocal microscopy

Mouse tissue labelled with FISH was imaged using a spinning disk Operetta CLS (Perkin Elmer). Sections were imaged in confocal mode using a sCMOS camera and a 40x automated-water dispensing objective. Sections were imaged with z-stacks at intervals of 1 μm. ROIs included the ME and CC. Gain and laser power settings remained the same between experimental and control conditions within each experiment.

### Light microscopy

Toluidine blue-labelled tissue was imaged using a (Nikon Eclipse E600) light microscope with 40x and 100x (dry) objectives. The ROI was the ME.

### Spinning disk confocal microscopy

Thick cleared mouse tissue was imaged using an Andor Dragonfly spinning disk confocal with a 20x objective. Sections were placed in a glass-bottom dish (MatTek – P-35G-0-14-C) with a small amount of 1:1 oil mixture to coat the interface of the glass and tissue. Sections were imaged with z-stacks at the software recommended interval. The ROI was the ME and ARC – the large field of view of this microscope allowed both structures to be imaged at once without tiling.

## Image analysis

For all experiments, the images were blinded to experimental condition before quantification.

### Mouse FISH

Harmony software (Perkin Elmer) was used to automatically quantify number of labelled RNA molecules (spots) per cell, intensities of spots, and area of spots.

### Immunohistochemistry, human FISH, toluidine blue labelling

For immunolabelled, toluidine blue labelled mouse sections and FISH labelled human sections, Fiji software was used to analyse colocalization, distribution, and counts/density of markers^46^. For images in z-stacks, individual images were first projected into a single image (a ‘Z-Project’ at maximum intensity) so all cells could be counted at once, and to eliminate double-counting. Areas of ROIs were measured by setting the image scale according to scale bars imprinted on images during acquisition, then tracing the ROI with the freehand tool and measuring. Borders of the ME, ARC, and other ROIs were determined using the Paxinos and Franklin Mouse Brain Atlas^39^. The Fiji manual cell counter was used to count marker-positive cells or axons. To quantify cell size, 10 cells per section were measured by tracing their perimeters with the Fiji freehand tool.

### Synaptic puncta quantification

Stacks of microphotographs from confocal microscopy were analyzed with Imaris software (Bitplane). Surface rendering was done for each channel prior to contact analysis. Contacts between synapses and OPC somas and elongations were determined using the “Surface Surface Contact Area” plug-in (Imaris, Bitplane). Numbers of contacts between VGAT-positive elements and VGluT1-positive elements were counted on each PDGFR α-positive elements of OPC cells. Scores were standardized by calculating the ratio between VGluT1-positive elements and VGAT-positive elements per OPC cell.

### TNR and WFA staining quantification

20um stacks taken at 1um intervals were analysed using Imaris software (Bitplane). Surfaces were reconstructed in 3D for each staining. Surface, volume and mean intensity were exported for each TNR+ and WFA+ object.

### Cleared tissue visualization

Videos of thick cleared tissue were made using Imaris software (Oxford Instruments).

### Myelin thickness

To assess differences in g-ratio between fasted and refed conditions, at least 100 distinct transverse axons were measured per animal. Using Fiji, the cross-sectional area of axons was measured by tracing the outside of the axon with the freehand tool. Similarly, the outside of the myelin sheath was traced to determine the area of the myelin+axon. Diameters of the axon and the myelin+axon were back-calculated from the areas, assuming the cross sections were perfect circles. The g-ratio was then calculated by dividing the axon diameter by the myelin+axon diameter.

## Statistical analysis and data visualization

All code used for analysing data and creating figures related to scRNAseq is found in the online repository Github: Kohnke-et-al-2019. Ranking of cluster-defining genes and statistical significance of differentially expressed genes (DEGs) was determined by the cellrangerRkit (10x) and edgeR^35,36^ packages in R. Statistical significance of pathways changed and upstream regulators of DEGs between fasting and refeeding was determined using Ingenuity Pathway Analysis (Qiagen). Figures relating to scRNAseq data were created using the cellrangerRkit, ggplot2, tidyr, and GOplot packages^47–50^. For histology experiments, GraphPad Prism 8 (GraphPad Software) was used to define statistical significance and create graphs. Two-sided unpaired Student’s t-tests were used to compare data between fasted and refed conditions (averages per animal) and ANCOVA was used to compare slopes of linear regression lines. All data are expressed as mean ± SEM.

