## Supplementary material for "Nutritional signals rapidly activate oligodendrocyte differentiation in the adult hypothalamic median eminence": Merged pdf od supplementary figures

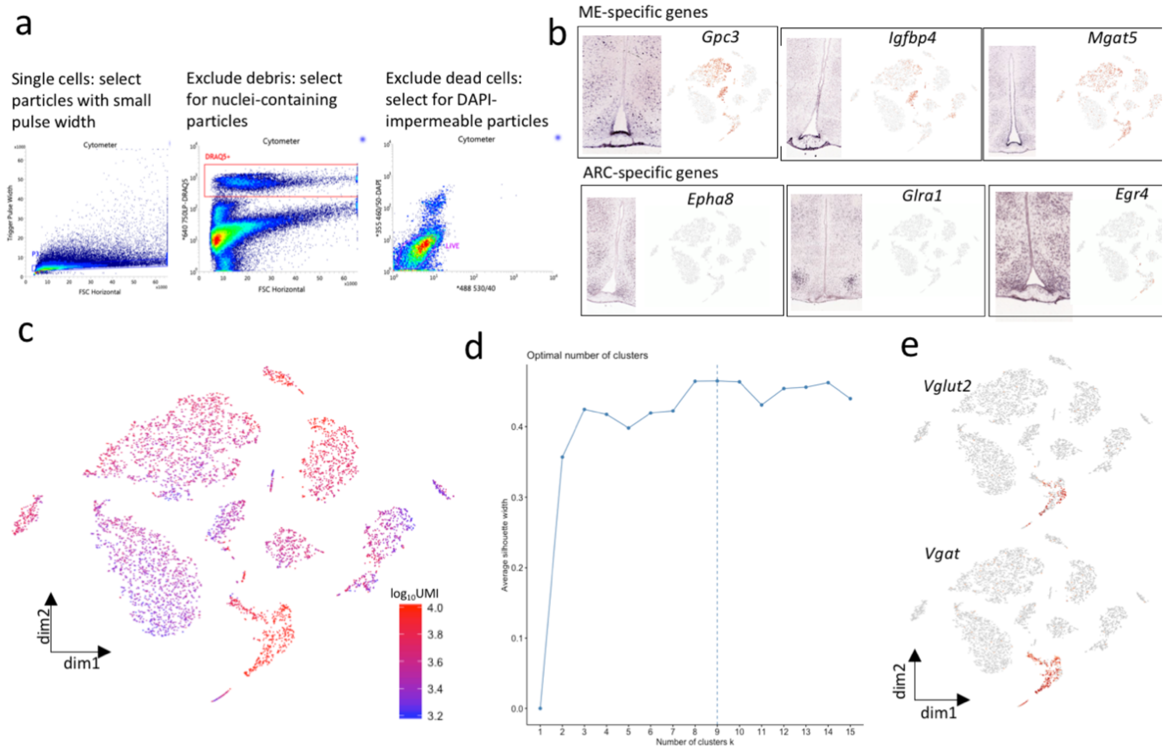


**Supplementary Figure 1.** (a) FACS cell sorting strategy (b) Allen Mouse Brain ISH gene expression images compared to gene expression in current dataset for ME / ARC specific genes (red labels high expression of the gene in the tSNE plot) (c) $Log_{10}UMI$ counts per cell mapped on tSNE plot (d) Silhouette analysis (optimal cluster analysis) of tSNE object coordinates (e) GABAergic (express vesicular GABA transporter, Vgat) and glutamatergic (express vesicular glutamate transporter 2, Vglut2) gene expression in current dataset

a

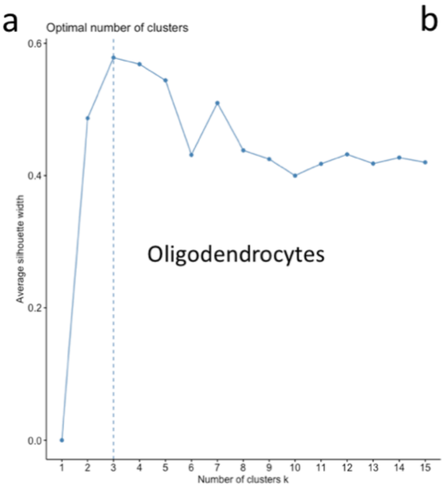


b

| Present study | | Campbell et al 2017 | | Marques et al 2016 | |
|---|---|---|---|---|---|
| OL name | markers | OL name | markers | OL name | markers |
| OPC | Cspg5, Pdgfra | OPC | Cspg4 | OPC | Pdgfra, Cspg4 |
| NFOL | Bmp4, Fyn, Tcf7l2 | early stage OLC | Mag, Bmp4 | COP | Bmp4, Fyn, Gpr17 |
| | | | | NFOL1-2 | Tcf7l2, Casr |
| MOL | Mal, Aspa | mid stage OLC | Mag, Man1a | MFOL1-2 | Mal, Mog, Plp1, Opalin |
| | | late stage OLC | Gm21984, Mag | MOL1-6 | Klk6, Pmp22 |

c

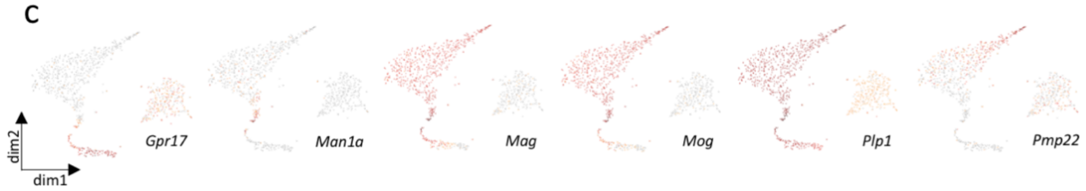


**Supplementary Figure 2.** (a) Silhouette analysis (optimal cluster analysis) of tSNE object coordinates (b) Comparison of terminology and defining markers between current study and two others examining single-cell transcriptomes of OLs (c) tSNE plots of current dataset showing expression of defining OL markers used in other studies. Red = high expression, grey = low expression

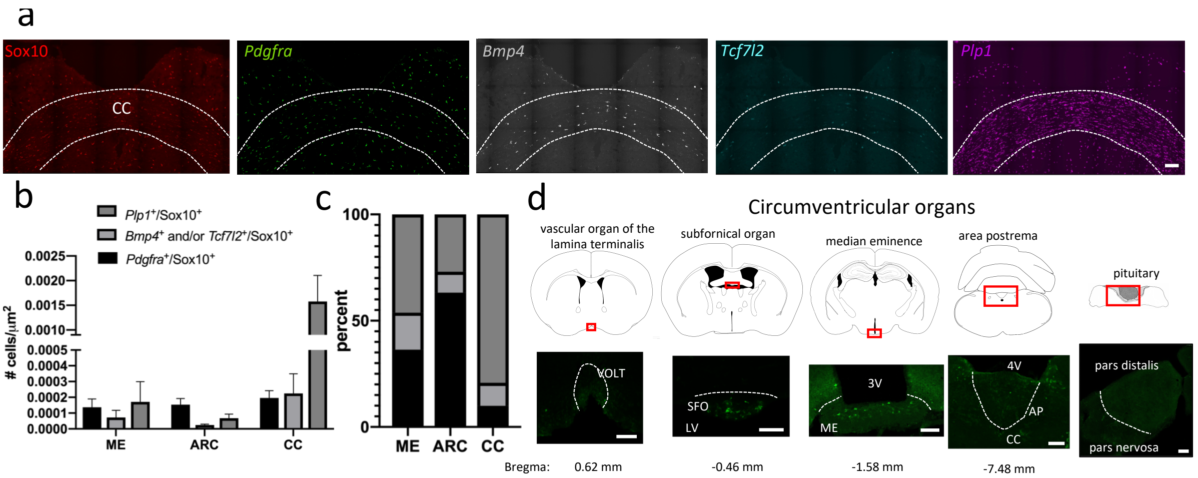


**Supplementary Figure 3.** (a) RNAscope combined to immunohistochemistry enables detection of OL subtype marker gene expression in the corpus callosum: red = Sox10 (IHC, pan-OL marker), green = *Pdgfra* (OPC marker), grey = *Bmp4* (NFOL marker), blue = *Tcf7l2* (NFOL marker), purple = *Plp1* (MOL marker). Scale bar = 100 µm (b) The ME, ARC, and CC have similar densities of *Pdgfra*$^+$/Sox10$^+$ cells (OPCs) but *Bmp4*$^+$ and/or *Tcf7l2*$^+$/Sox10$^+$ cell (NFOL) density is highest in the ME. The CC (a white matter tract) has the greatest density of *Plp1*$^+$/Sox10$^+$ cells (MOLs). Error bars depict mean ± SEM (c) Proportions of OPCs/NFOLs/MOLs differ between the ME, ARC, and CC (d) APC (green, a postmitotic OL marker) labels cells in the SFO, ME, and AP but not in the VOLT or pituitary. Scale bars = 100 µm

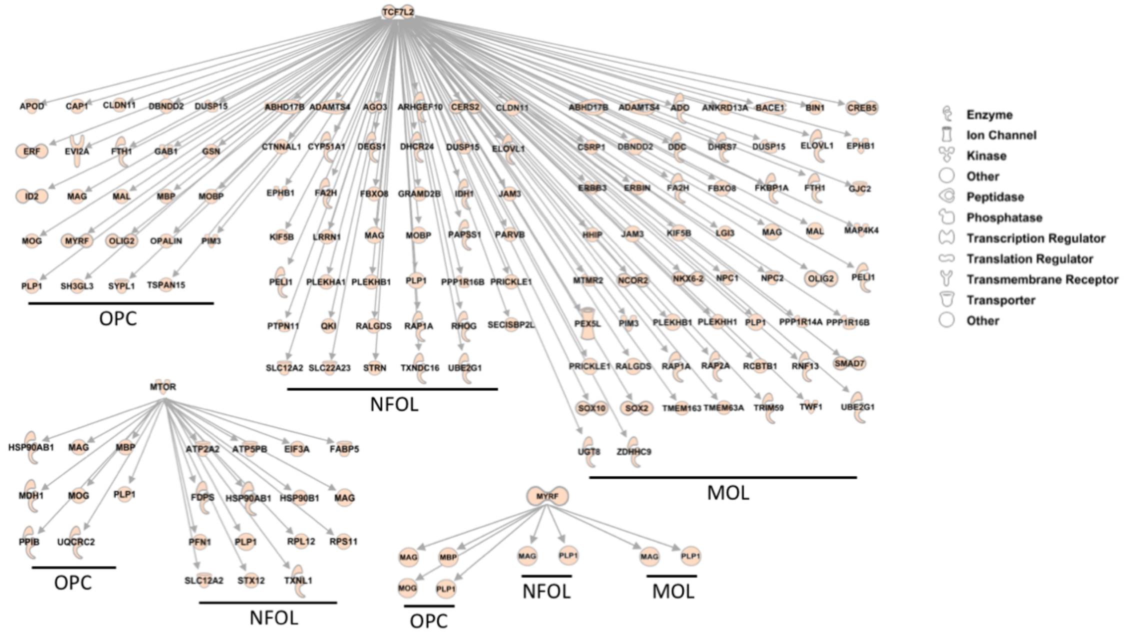


**Supplementary Figure 4**. Genes downstream from top upstream regulators Tcf7l2*,* Mtor, and Myrf

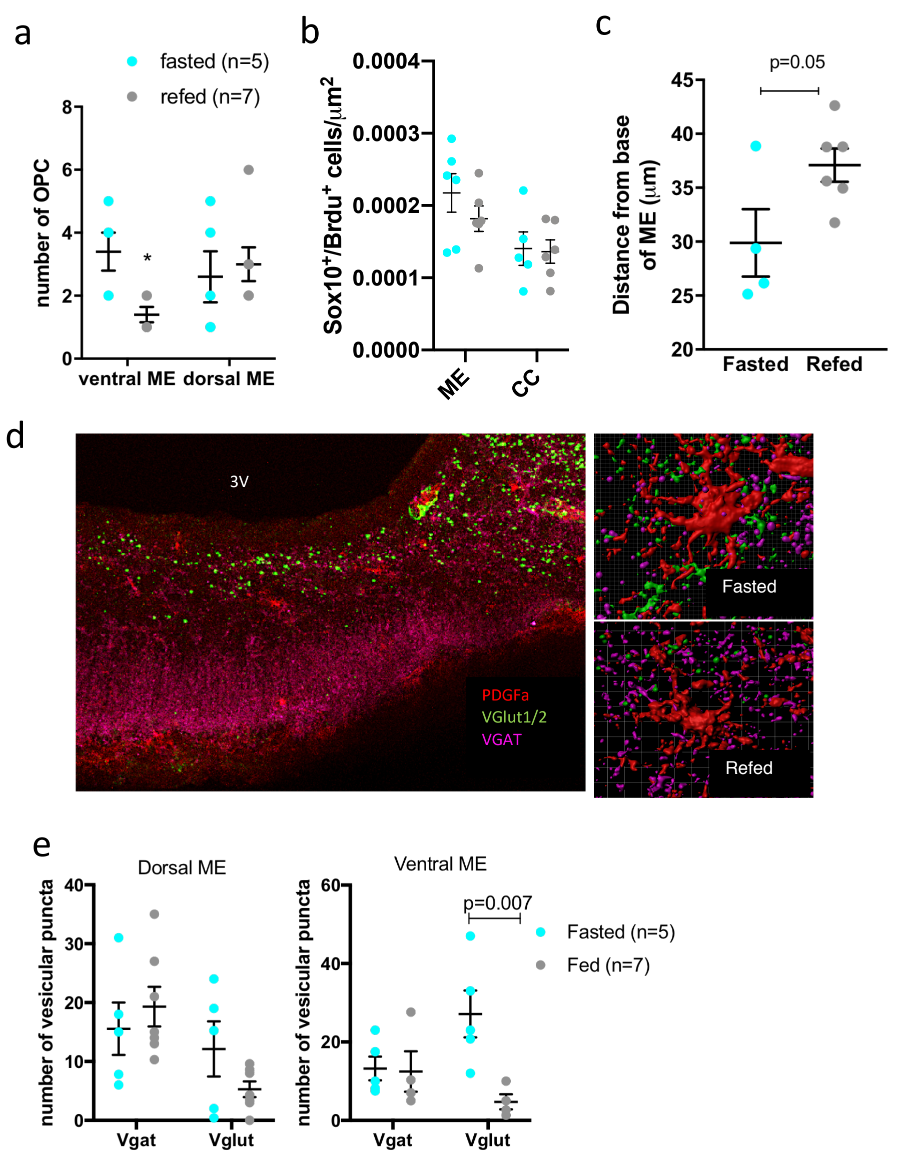


**Supplementary Figure 5. Refeeding promotes OPC differentiation and migration to the dorsal ME.** (a) Number of OPCs (PDGFRa+ cells) in the ME of mice fasted overnight (turquoise) or fasted overnight and refed for 1h (grey). (b) BrdU incorporation during a 24h fast (turquoise) or a 24h fast followed by a 1h refeed in the ME and CC. (c) Dorsoventral localization of NFOLs (Brdu$^+$, Sox10$^+$, Pdgfra$^-$ cells) in the ME of mice fasted overnight (turquoise) or fasted overnight and refed for 1h (grey). (d) Immunodetection of OPCs (PDGRA$^+$), glutamatergic (Vglut1/2$^+$) and GABAergic (VGAT$^+$) puncta and 3D surface reconstruction with Imaris. (e) Quantification of glutamatergic and GABAergic puncta contacting ME OPCs in mice fasted overnight (turquoise) or fasted overnight and refed for 1h (grey). Data are means ± SEM

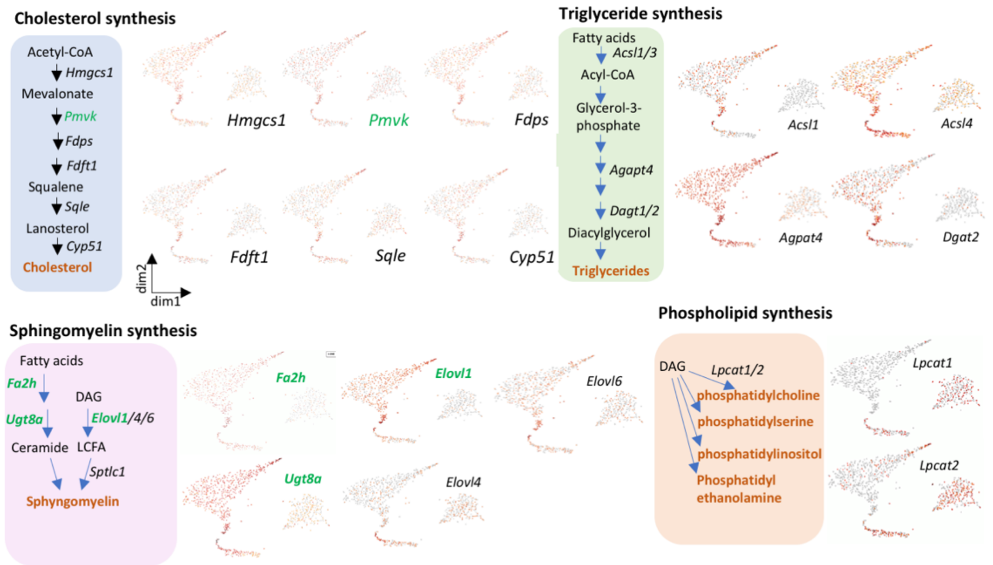


**Supplementary Figure 6.** Expression of genes encoding for proteins involved in cholesterol synthesis, triglyceride synthesis, sphingomyelin synthesis, and phospholipid synthesis in OPCs, NFOLs and MOLs clusters. Gene names in green are significantly upregulated upon 1h refeeding. Red = high expression, grey = low expression

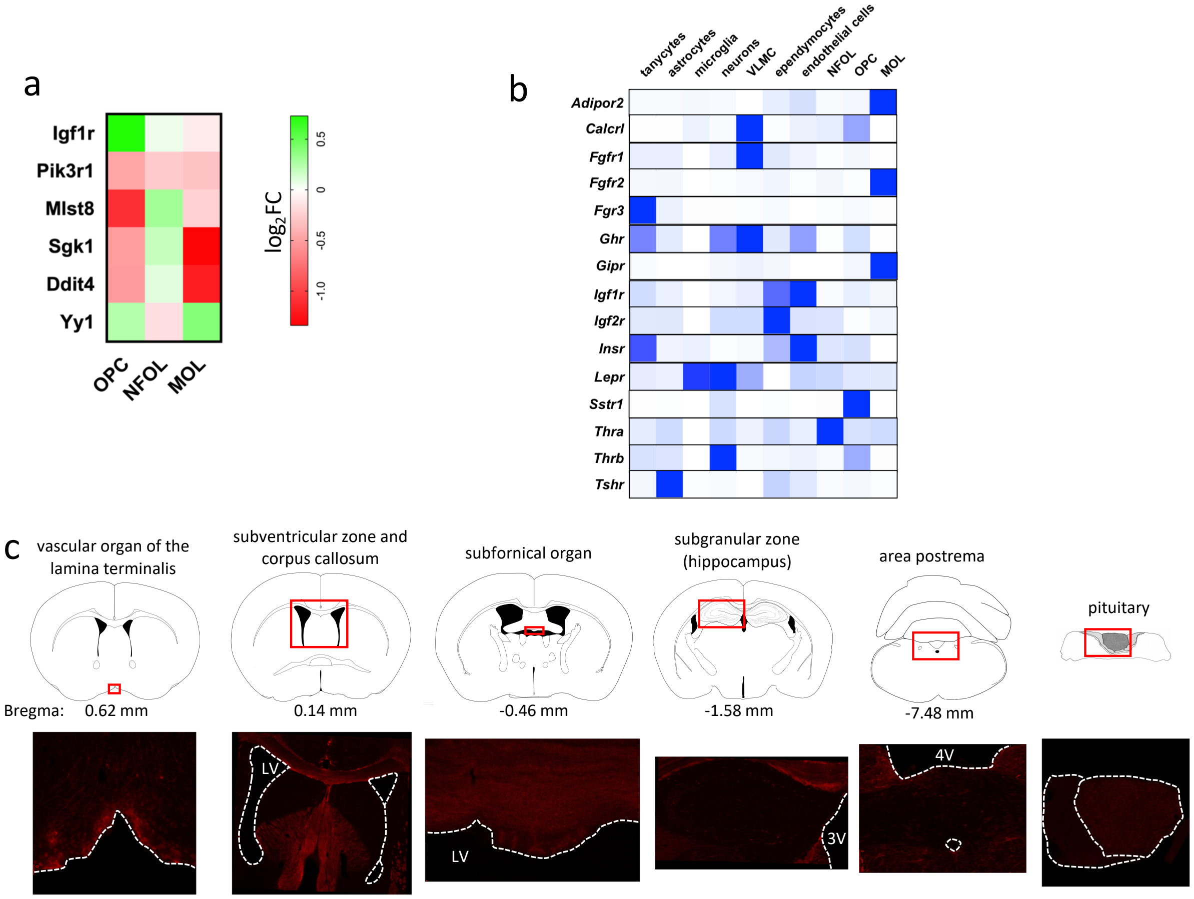


**Supplementary Figure 7.** (a) Heatmap of genes involved in mTOR signaling that are differentially expressed in scRNAseq dataset ($p < 0.5$, FDR < 0.25, *Pik3r* produces the p85 protein). (b) Relative expression of receptors for metabolic hormones in various cell types present in the ME. Scale is variable. (c) Labelling with the antibody to pmTOR (Ser 2448) in white matter tracts and other circumventricular organs throughout the mouse brain.
